# Macrophages drive the earliest anti-tumoral response to BCG therapy by directly killing bladder cancer through TNF signaling

**DOI:** 10.1101/2024.01.05.574391

**Authors:** Mayra Martinez-Lopez, Cátia Rebelo de Almeida, Marcia Fontes, Raquel Valente Mendes, Stefan H.E. Kaufmann, Rita Fior

## Abstract

The Bacillus Calmette-Guérin (BCG) vaccine is the cancer immunotherapy longest in use. Despite its effectiveness in bladder cancer (BC), its initial mechanisms of action remain largely unknown. Therefore, proper diagnostic assessments to identify patients who will not respond to treatment or develop resistance are lacking. Here, we set-out to unravel the earliest innate cellular mechanisms involved in BCG-induced clearance of tumors. We show that BCG induces a massive recruitment of macrophages to the tumor microenvironment and modulates their morphology and behavior towards a proinflammatory phenotype, while also promoting macrophage fusion-like events. We demonstrate that macrophages directly induce apoptosis and clearance of cancer cells through TNF-signaling and that they are indispensable for this antitumoral response since their depletion completely abrogates the BCG-anti tumor effect. Contrary to the general concept that macrophage antitumoral activities uniquely rely on stimulating an effective adaptive response, we demonstrate that macrophages alone can directly induce tumor killing and clearance; revealing an additional step to the BCG-induced tumor immunity model, that was not previously considered. In addition, we also provide proof-of-concept experiments demonstrating the potential of this unique *in vivo* preclinical model to test new innate immunomodulators.

## INTRODUCTION

The Bacille Calmette-Guérin (BCG) vaccine, based on the “Coley’s toxins” principle, is the cancer immunotherapy longest in use^1–3^. In bladder cancer, BCG is the most effective treatment to avoid disease relapse. Tumors staged as intermediate or high-risk non-muscle-invasive bladder cancer (NMIBC) are treated with intravesical BCG immunotherapy ∼two weeks after transurethral resection of bladder tumor (TURBT). BCG induction therapy consists of 6 weekly instillations after which maintenance therapy of 1 to 3 years is highly recommended to prevent progression and recurrence^4,5^. Despite being the gold-standard of treatment of NMIBC for 40 years, BCG intravesical immunotherapy has a high rate of adverse effects, there are worldwide shortages in its supply chain and some patients are resistant to treatment^2,6,7^. Additionally, the mechanisms through which BCG induces anti-tumor activity are not fully understood and BCG therapy has remained mostly unchanged^1,7,8^. Several studies have underscored the importance of a local inflammatory reaction in the bladder and a strong activation of the innate and adaptive immune systems upon BCG instillation^1,2,7,9^. The initial steps following instillation (∼120 minutes into the beginning of the treatment) have been elucidated through *in vitro* and murine studies and not all data has been supported by human studies.

A multi-step model of BCG-induced tumor immunity has been proposed^2^. Steps 1,2): upon treatment, BCG binds to and invades the bladder lumen, interacting with the urothelium and tissue-resident macrophages. Step 3): BCG is then internalized by immune cells, notably phagocytes^2,7,9^ and induces an innate immune response that triggers a strong local induction of pro-inflammatory cytokines and chemokines. This stimulates the recruitment of immune cell including neutrophils, monocytes, macrophages, T lymphocytes, B lymphocytes, and natural killer (NK) cells. Macrophages and other antigen-presenting-cells (APCs) present BCG antigens to T lymphocytes through the major histocompatibility complex (MHC) class II and trigger an adaptive immune response. Step 4): Therapy is thought to be successful if the induction of the adaptive immune response is biased towards Th1 cells^2,7^. The recruitment of all these immune cells leads to the development of granulomatous lesions in the bladder wall^7,9,10^.

Due to difficulties in assessing treatment response in patients, animal models of bladder cancer have been used to understand the mechanisms of BCG immunotherapy^11^. Historically, mice have been considered the gold-standard xenograft model due to their highly conserved genetic likeness with humans^12,13^. Nevertheless, the mouse xenograft model carries some disadvantages such as: the need for immunosuppressed or humanized animals and a large abundance of donor tumor cells (not compatible with biopsies or limited number of samples); long waiting times; high husbandry costs; moderate-low percentage of success in clinical trials. Additionally, single-cell live imaging is difficult due to their anatomical characteristics, namely skin and fur^13–16^.

The similarities in molecular pathways and drug responses between zebrafish and humans, and the ease in genetic manipulation have allowed for the development of robust cancer models. In zebrafish cancer xenografts, where human tumor cells are injected into zebrafish embryos or adults, cancer features such as proliferation, angiogenesis, metastasis and interactions in the tumor microenvironment (TME) can be rapidly visualized in real-time and at single cell level due to the optical transparency of the model^14,15,17–25^. Zebrafish xenografts have helped to elucidate the different chemo- and radiosensitivity profiles of several cancer cell types, highlighting their importance in future personalized medicine^22,26–29,30^. The role of the innate immune system in colorectal cancer progression and response to therapy has also been shown in this model^23,31^ and live imaging has allowed dissection of the earliest stages of cancer development and metastatic spread ^21,32–35^. Altogether, research in zebrafish cancer xenografts facilitates the rapid identification of novel cancer mechanisms that can be targeted by specific therapeutic approaches. In parallel, the zebrafish model has proven to be a powerful tool to study human tuberculosis (TB)^36–44^, in particular for the initial mechanisms involved in the pathophysiology of TB and granuloma development ^36–38,40–43,45^. This highlights the importance of the zebrafish model to study the role of the innate immune system in the development of complex pathologies.

Here, we characterized part of the initial innate immune response mechanisms that occur within the TME upon BCG treatment. We demonstrate *in vivo* in a bladder cancer zebrafish xenograft through real-time single-cell resolution microscopy that BCG immunotherapy induces cancer-cell apoptosis and clearance of tumors through macrophages and TNF signaling. BCG stimulated a massive recruitment of macrophages that were polarized towards a TNFα positive proinflammatory phenotype. Through high-resolution live microscopy we revealed that the presence of BCG in the TME induced profound changes in macrophage morphology and in cell-cell interactions. Innate immune cells were crucial for the anti-tumor effects of BCG, since in their absence tumor clearance was halted. Importantly, we demonstrate the utility of our xenografts in a preclinical setting, testing the efficacy of a newly genetically modified BCG vaccine (VPM1002-*M. bovis* BCGΔ*ureC*::*hly*)^46,47^ versus the conventional BCG vaccine. This next generation BCG-based vaccine is currently undergoing three phase III efficacy trials against TB and has already shown promising effects against bladder cancer^48^.

In summary, we dissected the earliest mechanisms of BCG immunotherapy and unveiled an additional step to the BCG-induced tumor immunity model; an active role of macrophages in the induction of tumor clearance, that was not previously regarded. Additionally, we provide proof-of-concept experiments for the use of zebrafish embryo xenografts in the preclinical setting to test new medicines aimed at boosting the host’s innate immune response, highlighting this model’s potential to become an integral part of future immunotherapy research.

## RESULTS

### BCG vaccine induces bladder cancer clearance and apoptosis

We started by developing a xenograft bladder cancer model for BCG immunotherapy in zebrafish embryos. For this purpose, we chose two bladder cancer cell lines, one isolated from a primary tumor staged as high risk non-muscle invasive bladder cancer (NMIBC-RT112)^49^ and another isolated from a tumor staged as muscle-invasive bladder cancer (MIBC-J82)^50^.

To optimize the BGC immunotherapy protocol and aware of the limited worldwide supply of intravesical BCG, we made use of the BCG vials (OncoTICE®) used in bladder cancer patients at the Day Hospital of the Champalimaud Foundation’s Clinical Centre. We labeled the bacteria with a lipophilic dye to allow for their identification and prepared them for injection. To generate the bladder cancer zebrafish embryo xenografts, bladder cancer cells were fluorescently labeled with a lipophilic dye and injected into the perivitelline space (PVS) of 2 days post-fertilization (dpf) zebrafish embryos as previously described^22,51^ (**Supplementary Figure 1**). At 1 dpi, bladder cancer xenografts were treated with one dose of intratumoral BCG, followed by a booster injection at 3 dpi and analyzed the following day (**Figure 1A**). Control xenografts followed the same treatment protocol but received PBS injections instead of BCG (**Figure 1A**). During the first week of zebrafish development only innate immunity is active (adaptive immunity is only mature at 2-3 weeks)^52–54^. Since our xenograft assay was performed during this first week of development, it provides an ideal temporal separation to specifically analyze the immediate effects mediated by innate immunity in the presence of cancer cells as a response to BCG treatment.

**Figure 1.**
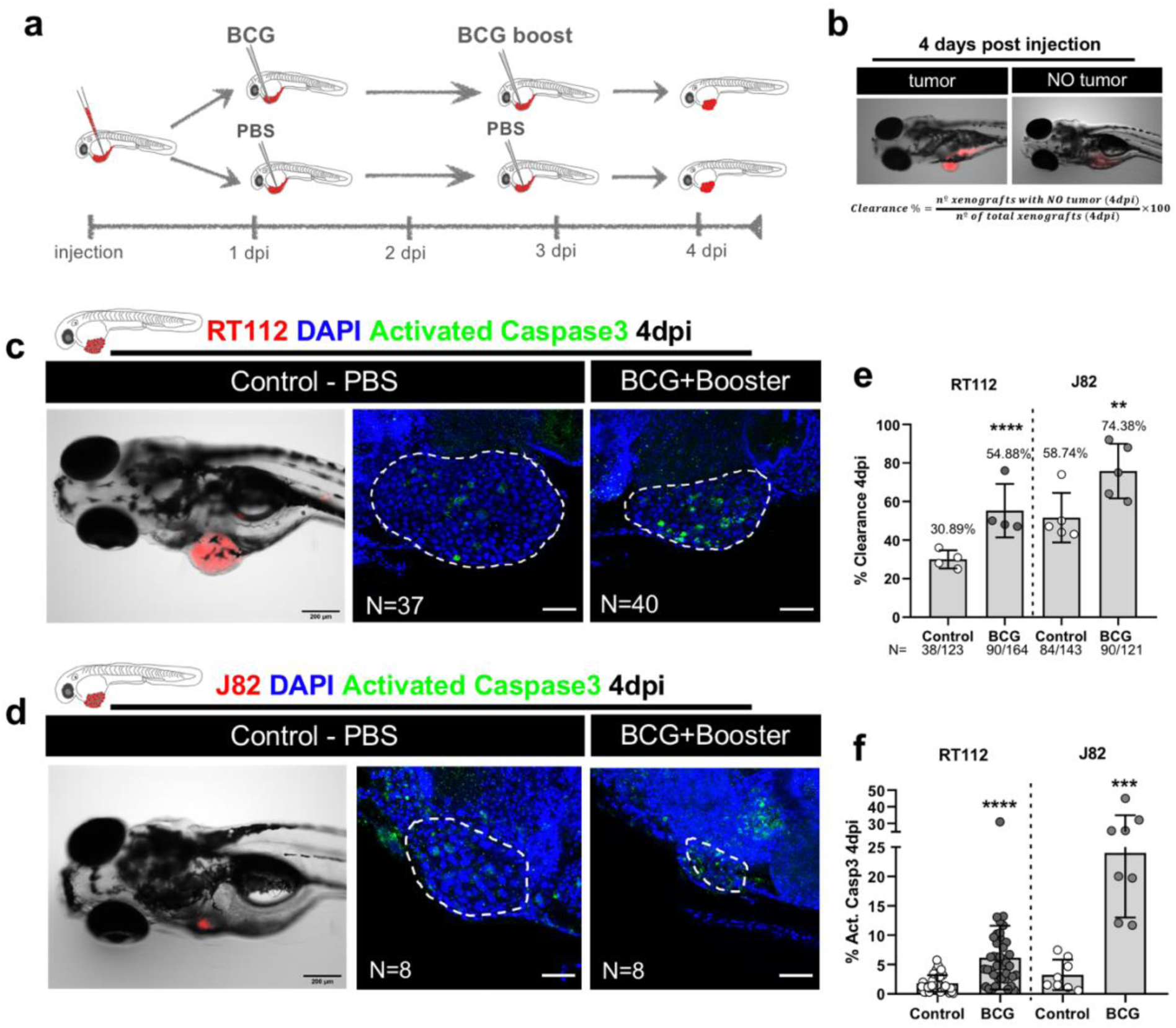
Zebrafish bladder cancer xenografts are susceptible to BCG therapy. **a)** Schematic representation of the BCG treatment protocol. **b)** calculation of clearance rate **c)** and **d)** Representative brightfield and confocal images of NMIBC-RT112 and MIBC-J82 control and BCG-treated xenografts with human cancer cells labelled in red, the apoptosis marker activated caspase 3 in green with DAPI nuclei counterstaining at 4dpi. Scale bar: 200µm. **e)** Quantification of the percentage of clearance in NMIBC-RT112 and MIBC-J82 xenografts at 4dpi. Bars indicate the results as AVG ± standard deviation of the mean (STDEV) and each dot represents a full round of injections in which N= # of xenografts without tumor at 4dpi/ total number of xenografts at 4dpi. **f)** Quantification of the percentage of apoptosis/activated caspase3 positive cells at 4dpi. Bars indicate the results as AVG ± STDEV and each dot represents one xenograft pooled from two independent experiments. White dashes outline the tumor. All images are anterior to the left, posterior to right, dorsal up and ventral down. Scale bar: 50 µm. dpi: days post-injection.

We assessed the impact of BCG treatment by evaluating *in vivo* tumor cell clearance, which is defined as the frequency of treated xenografts that lost the tumor mass at 4dpi. While both cell lines showed a baseline spontaneous tumor clearance/rejection, ∼30% in NMIBC-RT112 and ∼58% in MIBC-J82, BCG treatment almost doubled the clearance efficiency in NMIBC-RT112 xenografts (1.7 fold, ****P<0.0001). In MIBC-J82 xenografts, BCG also increased the efficiency of tumor clearance, but in a less pronounced manner (1.2 fold, **P=0.0076) (**Figure 1B-1D**). In conclusion, BCG efficiently induces bladder cancer cell clearance in the zebrafish embryo xenograft model.

The fact that BCG increased tumor clearance in the zebrafish xenografts raised the question of how human cancer cells were being cleared. We hypothesized that BCG could induce clearance either by direct cytotoxicity leading to cell death or by the stimulation of innate immune cells.

To tackle this question, we evaluated activated caspase 3, which marks cells undergoing apoptosis. We found that BCG treatment induced apoptosis of bladder cancer cells (NMIBC-RT112, ****P<0.0001; MIBC-J82, ***P=0.0002) (**Figure 1C-1E**), which suggested that BCG treatment was promoting an active clearance of cancer cells by the generation of programmed cell death. However, given that some bladder cancer cell lines are susceptible to direct toxicity induced by BCG *in vitro*^55^, the question remained whether this could be a direct consequence of BCG toxicity or an active process of cancer cell elimination mediated within the host TME. Thus, we determined whether BCG is toxic for NMIBC-RT112 and MIBC-J82 tumor cell lines *in vitro*. BCG treatment did not significantly affect the survival of cultured cancer cells. That is, vehicle and BCG-treated cells showed similar average cell numbers per field and similar abundance of apoptosis (**Supplementary Figure 2**). Thus, BCG is not directly toxic for NMIBC-RT112 and MIBC-J82 tumor cells, suggesting that the host TME is actively involved in tumor cell death.

### BCG induces infiltration of macrophages and polarization towards a pro-inflammatory phenotype

Since BCG treatment induced the elimination of human cancer cells in the zebrafish xenografts, we assumed that BCG modulates the innate response of the host embryo. Thus, to evaluate if BCG modulated the immune response and the TME, we quantified the presence of infiltrating neutrophils and macrophages, which are the main innate immune cells at this stage of zebrafish development^19^, in bladder cancer xenografts. To this end, we injected NMIBC-RT112 bladder cancer cells into *Tg(mpx:eGFP)*^56^ and *Tg(mpeg1:mcherry-F)*^56^ zebrafish hosts, in which neutrophils and macrophages are fluorescently labeled respectively (**Figure 2A**). We did not detect significant differences in the absolute numbers of infiltrating neutrophils between the control and BCG treated xenografts (**Figure 2B**). In contrast, we observed a significant increase in the absolute numbers of infiltrating macrophages upon BCG treatment (numeric doubling from a mean of 47 to a mean of 97, ****P=0.0001) (**Figure 2B**). Thus, these results indicate that BCG treatment induces macrophage recruitment into the TME.

**Figure 2.**
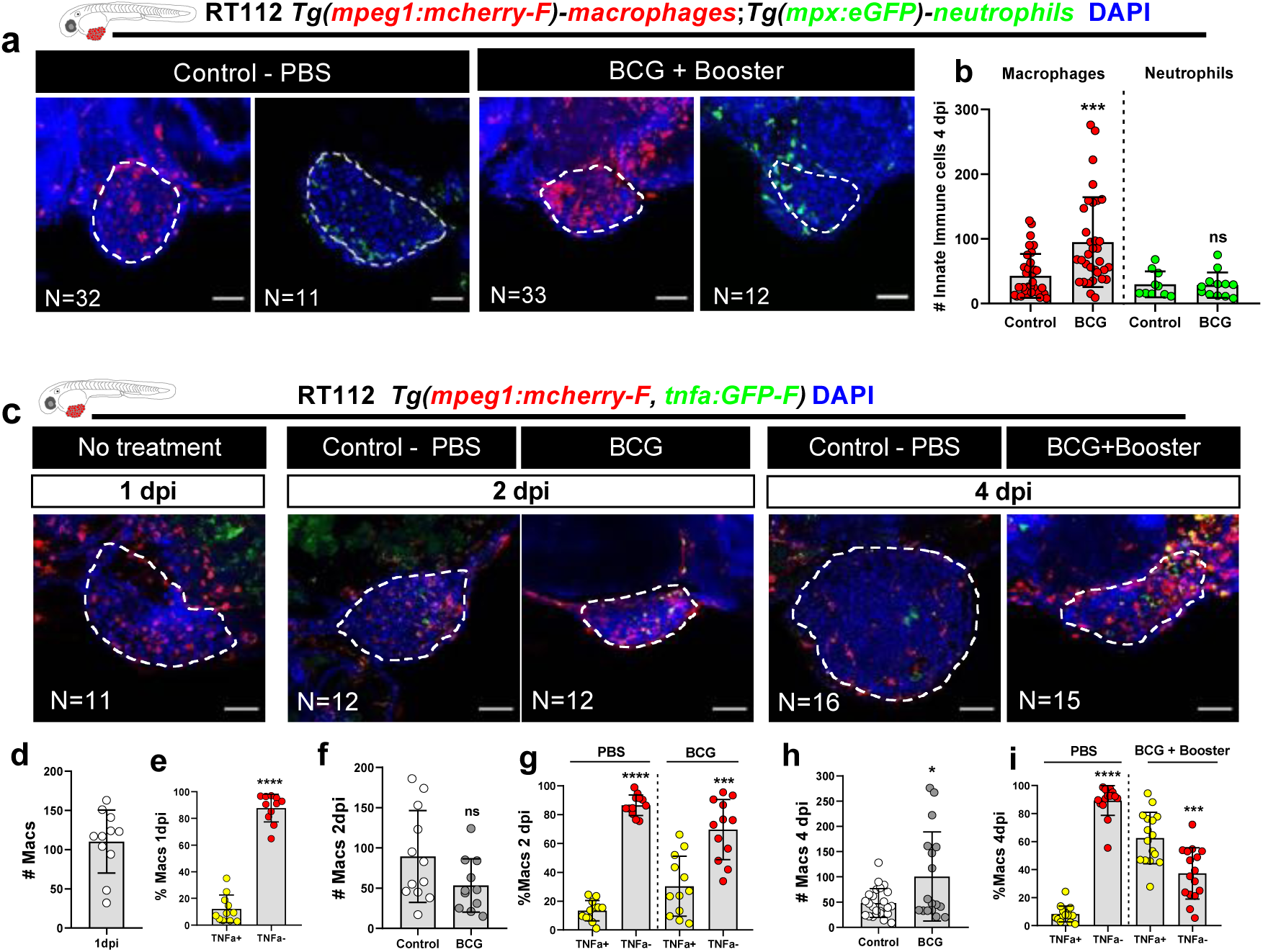
BCG modulates recruitment and polarization of macrophages in zebrafish bladder cancer xenografts. **a)** Representative confocal images of macrophages (red) and neutrophils (green) in NMIBC-RT112 control and BCG-treated xenografts. **b)** Quantification of the absolute numbers of infiltrating macrophages and neutrophils (***P=0.0003). **c)** Representative confocal images of TNFα expression (green) and macrophages (red) in NMIBC-RT112 control and BCG-treated xenografts. **d-i)** Quantification of the absolute number of macrophages and the percentage of TNFα positive and TNFα negative macrophages in the TME at 1dpi before treatment (****P<0.0001), 2dpi (***P=0.0001, ****P<0.0001), and 4dpi (***P=0.0001, **** P<0.0001). Bars indicate the results as AVG ± STDEV and each dot represents one xenograft pooled from 2 independent experiments. White dashes outline the tumor. All images are anterior to the left, posterior to right, dorsal up and ventral down. Scale bar: 50 µm. dpi: days post-injection.

Notably, while macrophage recruitment to the TME shows activation of the immune system by BCG, macrophage recruitment does not inform whether they contribute to the elimination of human cancer cells or not. This is because macrophages can adopt either a pro-inflammatory (M1-like) or anti-inflammatory (M2-like) phenotype with tumor suppressing or tumor promoting functions, respectively^57–60^. Thus, to investigate whether BCG modulates macrophage polarization towards a pro-inflammatory M1-like state, we analyzed the presence of TNFα producing macrophages, which are considered M1-like with tumor suppressing functions. For this, we generated bladder cancer xenografts in double transgenic zebrafish carrying a general macrophage mCherry reporter driven by the mpeg1 promoter and a GFP reporter driven by the TNFα promoter [*Tg*(*mpeg1:mCherry-F; tnfa:eGFP-F*)]^61^. Infiltrating macrophages were analyzed at 1dpi (**Figure 2C-2D-2E**), 2dpi (**Figure 2C-2F-2G**) and 4dpi (**Figure 2C-2H-2I**). Quantification of the immune cell populations revealed that at 1dpi, prior to treatment, macrophages were mostly TNFα negative. However, upon BCG treatment, macrophages gradually polarized towards a TNFα positive pro-inflammatory phenotype and, at 4dpi, TNFα positive macrophages represented ∼62% of the total macrophage population in the tumor of BCG treated xenografts; whereas in the controls they represented only ∼8% (**Figure 2C-2H-2I**; ****P<0.0001). In addition, BCG treatment also induced a change in macrophage morphology from a mesenchymal or dendritic-like morphology to an ameboid and vacuole-rich morphology (**Supplementary Figure 3**). These results suggest that tumor elimination driven by BCG treatment is mediated by proinflammatory macrophages with tumor suppressing activity.

### Macrophages mediate BCG-induced tumor clearance

BCG treatment activated an anti-tumor response by inducing clearance and apoptosis with a strong recruitment of macrophages and their polarization towards a TNFα-expressing M1-like phenotype in zebrafish xenografts, which suggested that macrophages play a critical role in this response. To test this notion, we pharmacologically depleted macrophages by using liposomes containing Clodronate (L-clodronate), which are selectively phagocytosed by macrophages. For this, we injected 0.02ug of L-clodronate intratumorally at the same timepoints described in Figure1A.

Quantification of macrophages confirmed that L-clodronate efficiently reduced the number of macrophages in the TME and, almost completely abrogated their local presence (**Figure 3A**-**3C**). Remarkably, the anti-tumor effects of BCG, namely induction of tumor clearance and apoptosis, were fully abolished upon macrophage depletion (**Figure 3B-3C-3D-3E**). The same phenotype was observed in MIBC-J82 xenografts (**Supplementary Figure 4**). Interestingly, when comparing the L-PBS controls to L-clodronate treated xenografts that did not receive BCG treatment, we observed that the depletion of macrophages resulted in a reduction of spontaneous clearance (**Figure 3**). We conclude that BC tumor cells are spontaneously eliminated by macrophages, and that BCG treatment profoundly elevates their tumor clearance activity.

**Figure 3.**
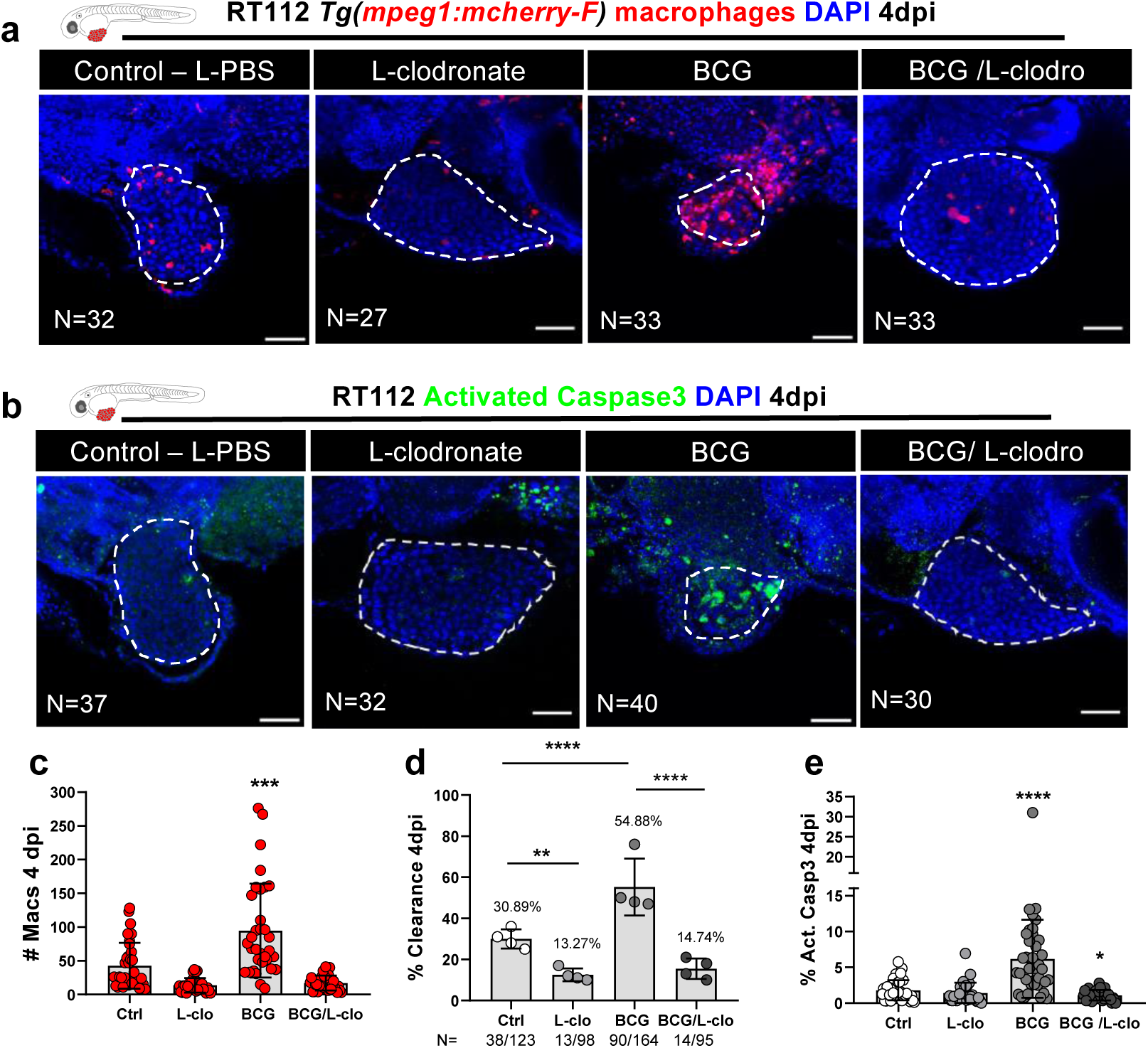
Macrophages are essential for susceptibility of zebrafish bladder cancer xenografts to BCG immunotherapy. **a)** Representative confocal images of infiltrating macrophages (red) in BCG/L-clodronate experiments. **b)** Representative confocal images of NMIBC-RT112 xenografts stained for the apoptosis marker activated caspase 3 (green) in BCG/L-clodronate experiments. **c)** Quantification of the absolute numbers of infiltrating macrophages in BCG/L-clodronate experiments (***P=0.0001). Bars indicate the results as AVG ± STDEV and each dot represents one xenograft pooled from 2 independent experiments. **d)** Quantification of the percentage of clearance in BCG/L-clodronate experiments at 4dpi (**P=0.0022, ****P<0.0001). Bars indicate the results as AVG ± standard deviation of the mean (STDEV) and each dot represents a full round of injections in which N= # of xenografts without tumor at 4dpi/ total number of xenografts at 4dpi. **e)** Quantification of the percentage of apoptosis/activated caspase3 positive cells in BCG/L-clodronate experiments at 4dpi (*P=0.0102, ****P<0.0001). Bars indicate the results as AVG ± STDEV and each dot represents one xenograft pooled from 3 independent experiments. White dashes outline the tumor. All images are anterior to the left, posterior to right, dorsal up and ventral down. Scale bar: 50 µm. dpi: days post-injection.

To further verify these results, and to rule out that the macrophage-dependent effect of BCG is a general feature of the zebrafish bladder cancer xenografts, we treated NMIBC-RT112 xenografts with the cytotoxic drug Mitomycin C^5^. As expected, Mitomycin C exerted its anti-tumor cytotoxic effect even in the absence of macrophages (**Supplementary Figure 5**). All together, these findings revealed that the initial tumor clearance and induction of apoptosis upon BCG immunotherapy is mediated by macrophages that are recruited to the bladder tumor. BCG’s mode of action in this model is through the innate immune system and not through direct BCG toxicity on the cancer cells.

### VPM1002 is more efficient in inducing tumor clearance and a pro-inflammatory tumor microenvironment than the conventional BCG vaccine

We next tested the tumor suppressing efficiency of the standard BCG vaccine in comparison with a novel promising next generation vaccine candidate of BCG, the VPM1002 vaccine (*M. bovis* BCGΔ*ureC*::*hly*)^1,2,47,62,63^.

VPM1002 is a genetically modified BCG vaccine strain. In this strain, the urease C encoding gene was replaced by the listeriolysin encoding gene. The listeriolysin gene is derived from *Listeria monocytogenes*, and its main role is to perturbate the phagosomal membrane provided the phagosomal milieu is acidic. This genetic modification confers the VPM1002 strain with higher immunogenicity by allowing mycobacterial antigens to escape to the cytosol of macrophages. Moreover, membrane perturbation allows egress of double-strand DNA which induces inflammasome activation resulting in generation of IL-1 and IL-18 as well as induction of LC32 as marker for autophagy/xenophagy^63^. VPM1002 is currently undergoing three phase III clinical efficacy trials to assess its efficacy in TB prevention in different populations in Sub-Saharan Africa and India^64–66^. A phase II clinical trial has also been performed to evaluate its effects in bladder cancer treatment in Switzerland and Germany ^67,46,68,63,69,47,48^.

Thus, we generated working stocks from live cultures of conventional BCG and VPM1002, and treated the bladder cancer xenografts by injecting BCG and VPM1002^48^ intratumorally. We followed the same treatment schedule shown in **Figure 1A**. We chose BCG:SSI as our control strain due to its genetic profile, which is closer to VPM1002^70^.

Our results show that both conventional BCG and VPM1002 strains were able to induce ∼45% of tumor clearance (**Figure 4A-4B**). However, and in alignment with previous studies^46,67^, VPM1002 induced a significantly higher infiltration of macrophages and a more pronounced tumor apoptosis in the TME than the conventional BCG vaccine (**Figure 4A-4C-4D-4E, \*\*\*\***P<0.0001). With regard to neutrophil infiltration, we could not detect significant changes among the two vaccines (**Supplementary Figure 6**).

**Figure 4.**
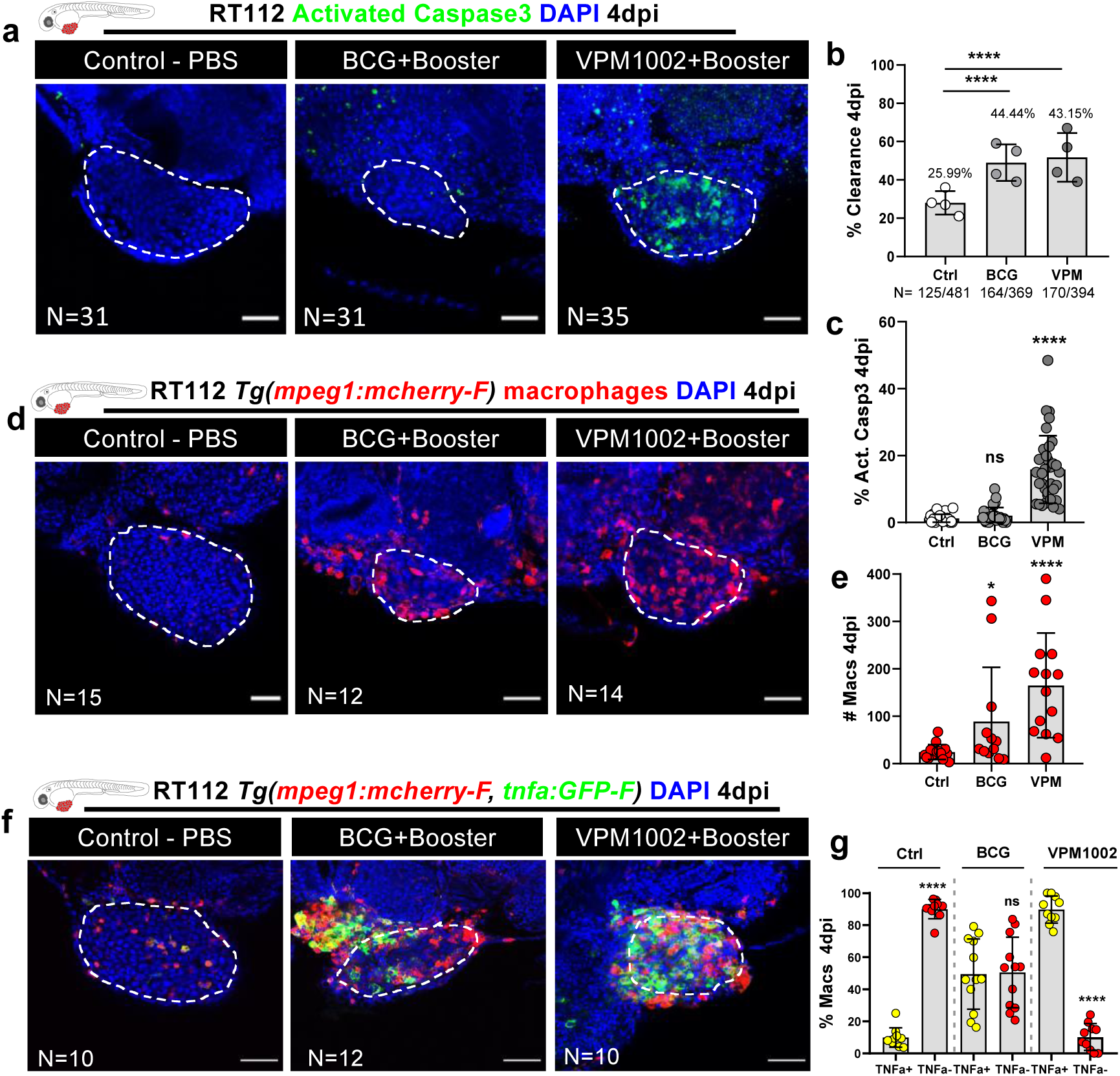
Zebrafish bladder cancer xenografts are susceptible to immunotherapy with the conventional and genetically modified BCG strains. **a)** Representative confocal images of NMIBC-RT112 control and BCG- or VPM1002-treated xenografts stained for apoptosis (activated caspase 3 in green) at 4dpi. **b)** Quantification of percentage of clearance in NMIBC-RT112 control and treated xenografts at 4dpi (****, P<0.0001). Bars indicate the results as AVG ± standard deviation of the mean (STDEV) and each dot represents a full round of injections in which N=# of xenografts without tumor at 4dpi/ total number of xenografts at 4dpi. **c)** Quantification of the percentage of apoptosis/activated caspase3 positive cells in NMIBC-RT112 control and treated xenografts at 4dpi (****, P<0.0001). **d)** Representative confocal images of infiltrating macrophages (red) in NMIBC-RT112 control and treated xenografts. **e)** Quantification of absolute numbers of infiltrating macrophages in NMIBC-RT112 control and treated xenografts at 4dpi (*P=0.0308, ****P<0.0001). **f)** Representative confocal images of TNFα expression (green) and macrophages (red) in NMIBC-RT112 control and treated xenografts. **g)** Quantification of the percentage of TNFα positive and TNFα negative macrophages in the TME of NMIBC-RT112 control and treated xenografts at 4dpi (****P<0.0001). Bars indicate the results as AVG ± STDEV and each dot represents one xenograft, from 3 independent experiments. White dashes outline the tumor. All images are anterior to the left, posterior to right, dorsal up and ventral down. Scale bar: 50 µm. dpi: days post-injection.

The conventional BCG vaccine polarized macrophages towards a pro-inflammatory phenotype at 4dpi (from ∼11% of TNFα positive macrophages in controls to ∼50% in BCG treated xenografts), but the VPM1002 vaccine was much more efficient in generating a highly pro-inflammatory TME with ∼90% of the macrophages being TNFα positive (****P<0.0001) (**Figure 4F-4G**). These results are in agreement with inflammatory phenotypes described in macrophages *in vitro* and in mice after VPM1002 exposure^46,67^.

### BCG and VPM1002 vaccines stimulate macrophage kinetics and their inter-cellular interactions

We next used light sheet imaging to further understand how macrophages respond to conventional BCG and VPM1002 vaccines and provide a real-time visualization with single cell resolution of these processes. At 1 dpi, immediately after treatment, control or BCG/VPM1002 treated *Tg(csf1ra:GFP)*^71^ xenografts, in which macrophages are fluorescently labelled in green, were imaged throughout 15 consecutive hours to assess the macrophage kinetics during this process (**Figure 5A; Video 1-3**).

**Figure 5.**
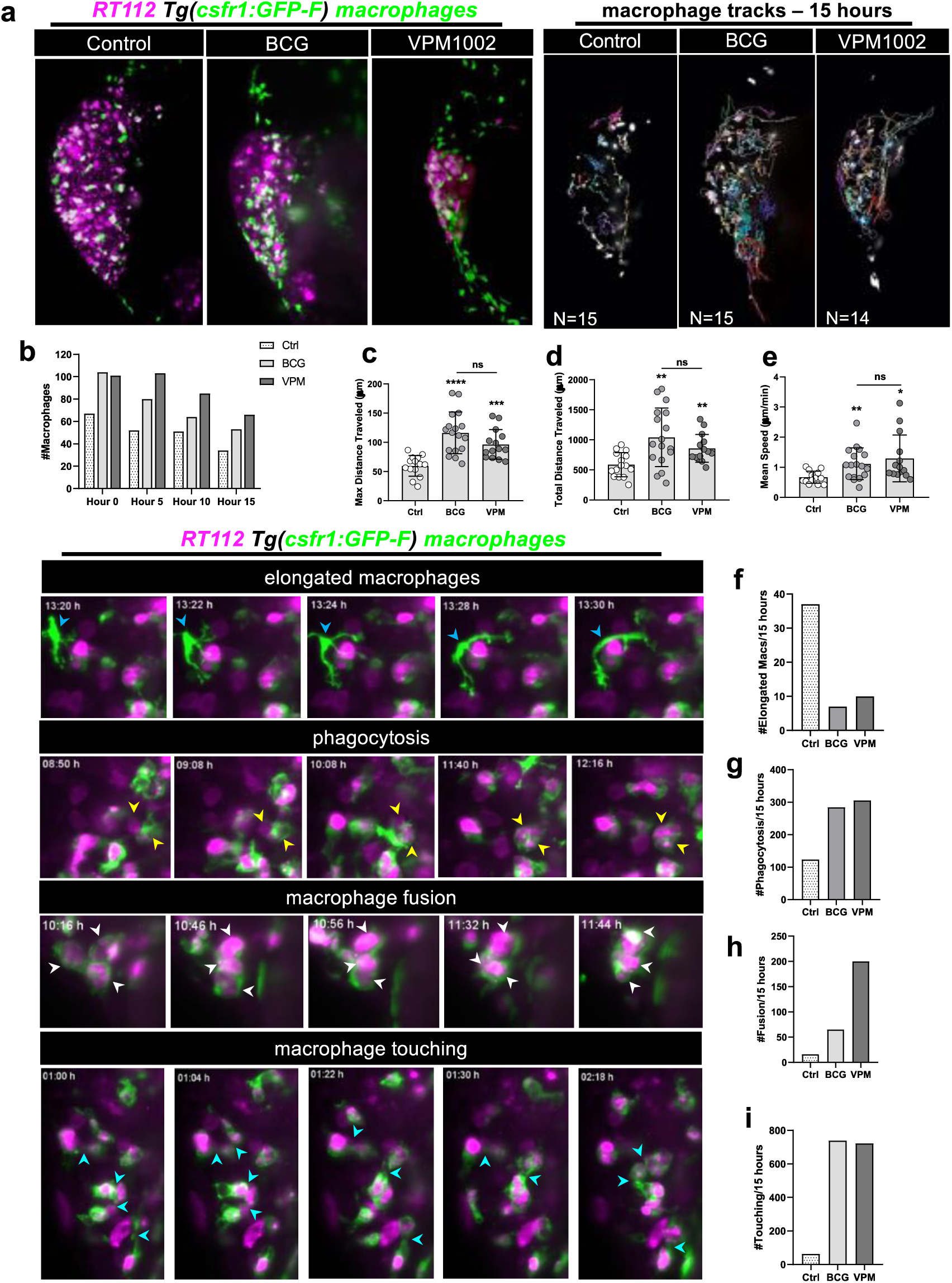
Live imaging reveals that BCG and VPM1002 vaccines stimulate macrophage kinetics and their inter-cellular interactions. **a)** Representative maximum intensity projections of cancer cells (magenta) and macrophages (green) at 10 hours of light sheet imaging. Representation of the macrophage tracks in which each colored line shows the path that an individual macrophage followed throughout 15 hours. **b)** Quantification of the absolute numbers of macrophages in NMIBC-RT112 control and BCG- or VPM 1002-treated xenografts at different timepoints during imaging. **c)** Quantification of the maximum distance travelled in microns (µm) by macrophages during 15 hours after treatment in NMIBC-RT112 xenografts (***P=0.0002, ****P<0.0001). **d)** Quantification of the total distance travelled in microns (µm) by macrophages during 15 hours after treatment in NMIBC-RT112 xenografts (BCG**P=0.0019, VPM1002 **P=0.0024). **e)** Quantification of the mean speed in microns (µm) per minute travelled by macrophages during 15 hours after treatment in NMIBC-RT112 xenografts (*P=0.0109, **P=0.0041). Representative still images of light sheet movies illustrating different macrophage - interaction events. Quantification of the number of elongated macrophages (blue arrow heads) **(f)**, number of phagocytic macrophages (yellow arrow heads) **(g)**, number of fusion events (white arrow heads) **(h)** and the number of membrane touching events **(i)** observed in 15 hours of imaging in NMIBC-RT112 xenografts. Bars indicate the results as AVG ± STDEV and each dot represents one macrophage.

Throughout the assay, the number of macrophages in the TME of the BCG and VPM1002 treated xenografts was higher than the control xenografts (**Figure 5B**). Quantification of the overall movement, distance-travelled and speed of macrophages revealed that these parameters were higher in both vaccine treated conditions compared to the control. That is, BCG/VPM1002 treatment induced changes in the behavior of macrophages and their interaction with surrounding macrophages. BCG/VPM1002 treatment not only increased cancer cell phagocytosis (**Figure 5G**), but also the frequency of touching (**Figure 5I; Video 4**) and fusion events (**Figure 5H; Video 5**) of macrophages. Interestingly, we noticed that elongated macrophages with no phagocytic capacity (dendritic-like) were more prevalent in control xenografts than in BCG/VPM1002 treated xenografts (**Figure 5F; Video 6**). Both vaccines induced similar macrophage behaviors, with VPM1002 inducing more fusion events than the conventional BCG (**Figure 5A-5C-5D-5E**). These fusion events are reminiscent of the initiation of granuloma-like structures.

Overall, these results show that the presence of BCG and VPM1002 in the TME generates an instantaneous mobile response in macrophages that migrate towards tumor cells. Phagocytic macrophages constantly and closely interact with each other. This process highlights the importance of cell-cell interactions in the BCG vaccine-mediated tumor clearance.

### BCG vaccine induces differentiation of L-plastin myeloid progenitors

It has been shown that BCG induces epigenetic changes in the hematopoietic compartment of human volunteers. These changes result in the skewing of hematopoietic stem cells towards myelopoiesis ^72^. Thus, we assessed whether we could also observe changes in the hematopoietic progenitors of the zebrafish xenografts upon BCG treatment. We started by quantifying the number of macrophages in the caudal-hematopoietic-tissue (CHT) at 2dpi and 4dpi (**Figure 6A**-**6B**), where hematopoiesis and myelopoiesis are actively occurring^73^. The absolute number of macrophages remained similar in control and BCG treated xenografts one day after treatment (**Figure 6B**). However, there was a significant increase in absolute macrophage numbers one day after the booster treatment in the CHT (**Figure 6A-6B**). Moreover, *in situ* hybridization for the hematopoietic markers *cmyb* (**Figure 6C-6D**) and *l-plastin* (**Figure 6E-6F**) suggested that BCG specifically stimulates myelopoiesis (*l-plastin*) and not general hematopoiesis (*cmyb*)^53,73^.

**Figure 6.**
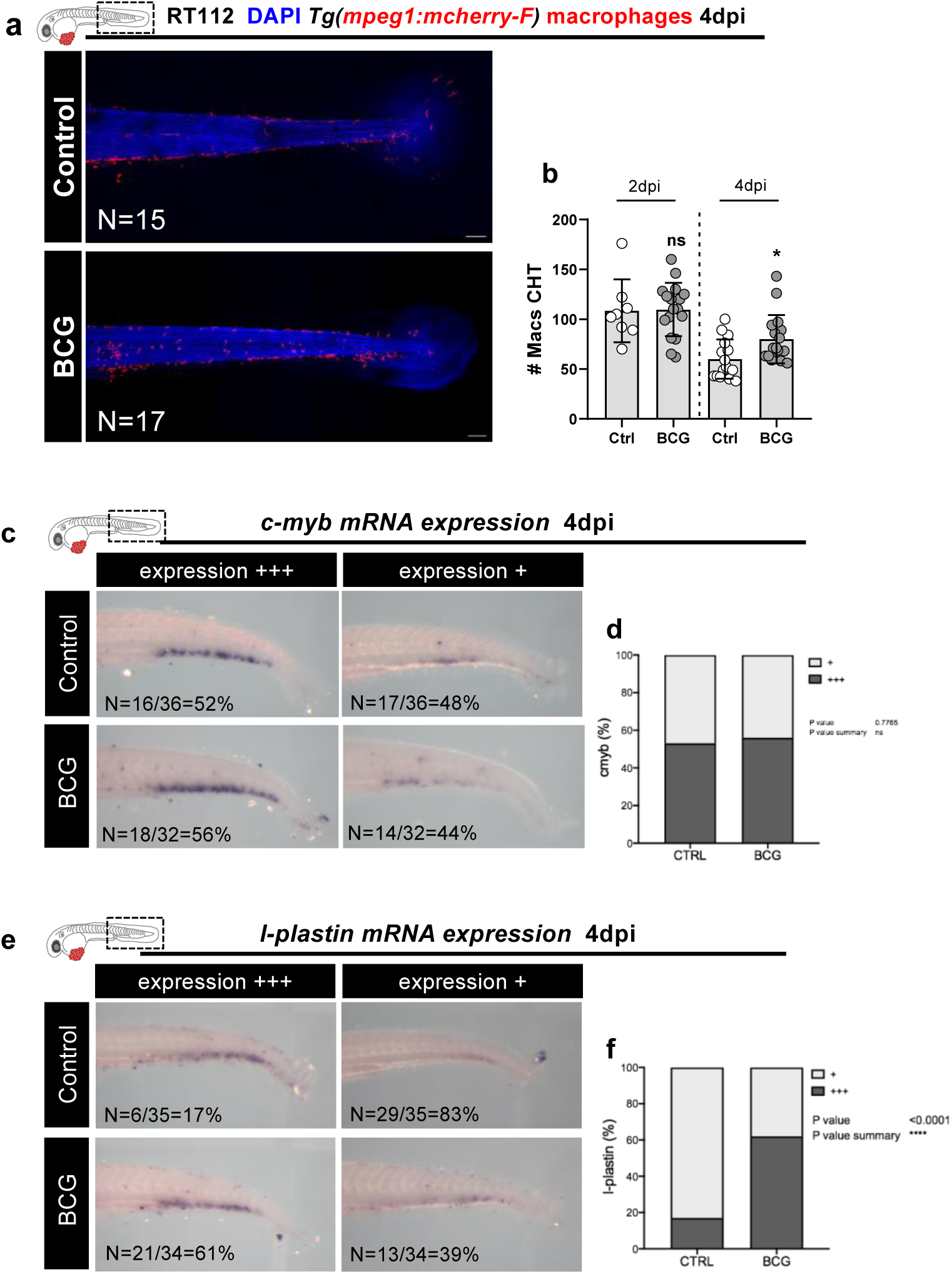
BCG induces myelopoiesis in zebrafish bladder cancer xenografts. **a)** Representative confocal images of macrophages (red) in the tails of NMIBC-RT112 control and BCG-treated xenografts at 4dpi. White dashes outline the CHT. **b)** Quantification of the absolute numbers of macrophages in the CHT at 2dpi and 4dpi (*P=0.0155). Bars indicate the results as AVG ± STDEV and each dot represents one xenograft. mRNA expression of *c-myb* (**c**) and l-plastin (**e**) and corresponding quantification (**d,f**, respectively). Number of analyzed xenografts are indicated in the figure. All images are anterior to the left, posterior to right, dorsal up and ventral down. Scale bar: 500 µm. CHT: Caudal hematopoietic tissue. dpi: days post-injection.

Then, we interrogated whether the strong innate immune response to BCG and VPM1002 immunotherapy was towards the bacteria alone or was dependent on the presence of bladder cancer cells. Thus, we challenged embryos without cancer cells to both vaccine strains and quantified the innate immune cells (**Supplementary Figure 8A**). Surprisingly, the absolute number of immune cells (macrophages and neutrophils) at 4dpi in the PVS of vaccine-treated embryos that were not carrying bladder cancer cells was similar to the PBS controls. Also, when we analyzed the whole-body distribution of the innate immune cells at 4dpi we could not see any significant differences between conditions (**Supplementary Figure 7A**). Along this line, the majority of macrophages in the vaccine treated embryos displayed a TNFα negative phenotype, similar to the control embryos (**Supplementary Figure 8A-8C**). Thus, these results indicate that the sole administration of BCG or VPM1002 does not trigger a sustained inflammatory response in the zebrafish embryos. In contrast, while injection of cancer cells alone already induced a mild recruitment of neutrophils and macrophages to the PVS (**Supplementary Figure 8B-8C-8D**), treatment with the conventional BCG (****P=<0.0001) and VPM1002 (****P<0.0001) vaccines induced a more profound recruitment of macrophages into the PVS region (**Supplementary Figure 8B-8D**). Altogether, these experiments suggest that a robust innate immune response requires both the presence of cancer cells and vaccine treatment to boost the infiltration and polarization towards a pro-inflammatory profile of macrophages in the TME, which then promotes tumor rejection.

### TNF**α**-signaling is essential for macrophage mediated anti-tumor activity

Our L-clodronate experiments, showed that the BCG anti-tumoral effect, clearance and apoptosis induction is macrophage dependent. However, at 4dpi, the tumors that escaped clearance showed a low percentage of infiltrating macrophages engaged in phagocytic behavior (∼26% of infiltrating macrophages in the controls, ∼10% in the BCG condition and ∼7% in the VPM1002 condition) regardless of the marked induction of apoptosis (**Supplementary Figure 9**). This led us to hypothesize that macrophages could induce cancer cell apoptosis through TNFα signaling, given the induction of TNFα expression in macrophages upon BCG treatment (**Figure 4F-4G**)

To test this, we employed the TNFα inhibitor Pentoxifylline (PTX)^74^ for treatment of xenografts in combination with the BCG/VPM1002 therapy or PBS in the controls (**Figure 7**). Our results show that inhibition of TNFα signaling completely abrogated the clearance process (**Figure 7D**), blocked the induction of apoptosis (**Figure 7E**), reduced macrophage recruitment (**Figure 7F**) and, as expected, also blocked the polarization of macrophages towards a proinflammatory phenotype^74^ (**Figure 7G**). Hence, TNFα plays a crucial role in successful BCG/VPM1002 treatment of bladder cancer.

**Figure 7.**
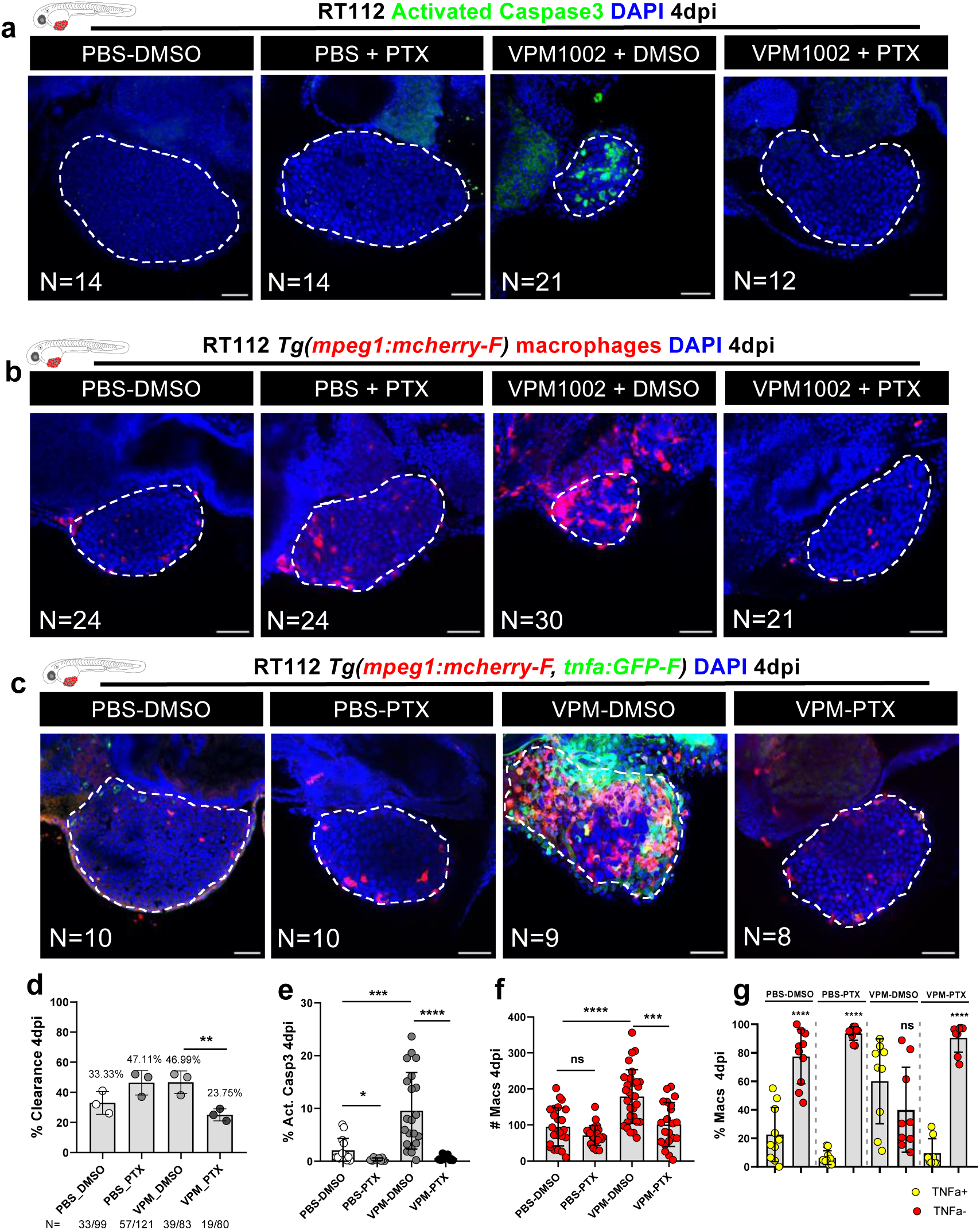
VPM1002 induction of bladder cancer cell clearance and apoptosis depends on TNFα signaling. **a)** Representative confocal images of NMIBC-RT112 control and VPM1002-treated xenografts exposed to either DMSO or PTX and stained for the apoptosis marker activated caspase 3 (green) at 4dpi. **b)** Representative confocal images of infiltrating macrophages (red) in NMIBC-RT112 control and VPM1002-treated xenografts exposed to either DMSO or PTX. **c)** Representative confocal images of TNFα expression (green) and macrophages (red) in NMIBC-control and VPM1002 treated xenografts exposed to either DMSO or PTX. Quantification of clearance (**d,** **P=0.0031), apoptosis/activated caspase3 (**e,** *P=0.0165, ***P=0.0002, ****P<0.0001), number infiltrating macrophages (**f,** ***P=0.0002, ****P<0.0001) and TNFα positive/negative macrophages in NMIBC-RT112 control and VPM1002-treated xenografts exposed to either DMSO or PTX at 4dpi (**g,** ****P<0.0001 ). Bars indicate results as AVG ± STDEV and each dot represents one xenograft pooled from 2 independent experiments. White dashed lines outline the tumor. All images are anterior to the left, posterior to right, dorsal up and ventral down. Scale bar: 500 µm. Scale bar: 50 µm. DMSO: dimethyl sulfoxide. PTX: pentoxifylline. dpi: days post-injection.

## DISCUSSION

The BCG vaccine was the first successful cancer immunotherapeutic agent. BCG elicits a non-specific immune response that promotes cancer clearance and prevents recurrence^75–77^. Despite its successful history, the precise mechanisms of action of BCG, in particular immediately after instillation, remain largely unknown^1,2,7–9,78^. In this work, we set to elucidate the initial anti-tumoral mechanisms of action of BCG through the use of the zebrafish bladder cancer xenograft model. For this, we focused on the cross-talk between BCG and innate immunity, which initiates the cascade of responses to therapy.

We showed *in vivo* that BCG induced tumor clearance and apoptosis of human bladder cancer cells and that this effect was mediated by macrophages. Immediately after BCG treatment, macrophages massively infiltrate tumors and become polarized towards a pro-inflammatory phenotype (M1-like, TNFα positive). Depletion of macrophages with L-clodronate completely abrogated the BCG anti-tumor effects, demonstrating that clearance and apoptosis are dependent on macrophage activity. Long-term light sheet microscopy revealed that macrophages altered their behavior in response to BCG, increasing phagocytosis, macrophage cell-cell interactions, and fusion events. Next, we showed that cancer cell clearance highly depends on TNF signaling. Importantly, expression of the myelopoietic progenitor transcription factor l-plastin was increased in the CHT upon BCG treatment, suggesting skewing of the hematopoietic compartment towards myelopoiesis. Moreover, we provide proof-of-concept experiments demonstrating that our model was able to discern distinctive innate immune responses to two different BCG vaccine strains. The conventional BCG and the recombinant second generation BCG-based vaccine VPM1002.

These findings provide key insights into the initial processes involved in BCG immunotherapy. We challenge the notion that macrophages are only APCs and secrete cytokines to induce an effective adaptive response. We show that, in contrast to what is shown in the current BCG-induced tumor immunity model^2^, macrophages are also able to directly induce apoptosis and clear cancer cells *in vivo*. This is in accordance with a previous report that indicates that macrophages can have direct anti-tumor activity *in vitro*^79^. In this work, authors show that macrophages and T lymphocytes can directly kill bladder cancer cells upon BCG stimulation, with T lymphocytes having a higher anti-tumoral activity. So far, we could not find any *in vivo* reports showing this direct active role of macrophages.

In all, our work suggests a new step to the multi-step model of BCG-induced tumor immunity (2): An earlier stage where macrophages are able to directly kill and clear tumor cells. Nevertheless, some cancer cells still escape (shown in our model by the few tumors that remained uncleared after BCG treatment). Then macrophages that are no longer able to kill and clear tumor cells, call forth the adaptive immune response through expression of cytokines, chemokines and antigen presentation, fully inducing a complete immune response to clear the remaining tumors cells.

Macrophages are innate immune cells with a unique transcriptional diversity and the capacity of switching their phenotype and function in response to diverse stimuli. Additionally, macrophages are crucial in the development of pathologies caused by different members of the genus mycobacterium (including BCG)^80^ such as TB and leprosy^81–84^. Therefore, we focused on deeper understanding the role of macrophages in the anti-tumoral effects of the BCG vaccine. Several studies have shown that the bladder cancer TME is highly immunosuppressive^85,86^. Anti-inflammatory macrophages (M2-like) being the main cellular subset found in histopathological samples of BCG-failed/resistant patients^87–89^. In accordance, we also observed that untreated bladder cancer xenografts had a TME enriched in anti-inflammatory (M2-like-TNFα negative) macrophages. However, upon BCG treatment there is an induction of an inflammatory TME together with tumor clearance and apoptosis, which is highly dependent on TNF signaling.

We revealed that the presence of the BCG vaccine in the TME was sufficient for immediately triggering a brisk change in macrophage dynamics. Macrophages were highly mobile in response to two different vaccine strains, the conventional BCG and VPM1002. However, those exposed to VPM1002 were more inflammatory and efficient at inducing tumor apoptosis. These results highlight the notion that not all immune cell infiltrates are similar and that further features should be analyzed to predict treatment response.

Despite the fact that we did not observe any differences in neutrophil infiltration at 4dpi, we do not discard the possibility of changes in neutrophil phenotypes upon BCG treatment at earlier or later time points in our assay.

Importantly, in the absence of cancer cells, BCG did not elicit a marked innate immune response in the zebrafish larvae at 4dpi, suggesting that it did not induce systemic inflammation. In line with this, in healthy human volunteers, intradermal BCG vaccination does not prompt a systemic inflammatory response^90,91^. Therefore, the local and cancer cell-specific response to BCG in our model could allow for the dissection of specific mechanisms that occur within the TME. These results are consistent with outcomes from clinical trials that underscore a reduced or completely absent systemic inflammation in patients that benefit from BCG vaccination for heterologous protection^92^.

Live imaging analysis showed that macrophages acquired different phenotypes in response to BCG. From the different phenotypes displayed, we identified fusion events among phagocytic macrophages in the xenografts that were treated with BCG. Fusion events were more prevalent in the VPM1002 treated xenografts. Here, phagocytic macrophages came in close contact and appeared to fuse with each other. These macrophages resembled granulomatous multinucleated giant cells (MGC). MGC formation is a macrophage-specific event that is highly evolutionarily conserved^93^. Although MGC function is not clearly defined, it is proposed that this event promotes more profound phagocytic and antimicrobial capacities^93^. Thus, we speculate that the macrophage fusion observed in long-term imaging experiments was the beginning of the formation of MGCs in early granuloma, supporting previous studies that revealed that the granulomas formation is an earlier event than as previously shown^94^.

Upon inhibition of TNF signaling, BCG failed to induce tumor clearance and apoptosis. TNFα is required for host protection against mycobacterial infections and for granuloma formation^95,96^. TNFα is a transmembrane protein that mediates cell-cell contact-dependent apoptosis. This process is achieved through the binding of TNFα to its receptor TNF-R1, which is generally highly expressed in cancer cells^97–99^. We speculate that BCG-induced contact-dependent macrophage killing also takes place in human cancer patients, since the abundance of TNF in the urine of bladder cancer patients is markedly increased after BCG instillation^100^. Consistently, macrophages of gastric cancer patients that received BCG immunotherapy expressed high levels of TNFα^101^.

Novel therapeutic approaches focused on the adaptive immune system are among the leading therapies for BCG resistant bladder cancer patients^102^. Unfortunately, when used as single agents, these therapies only benefit a small number of patients and carry numerous adverse events^103^. It has been suggested that several of these therapies fail due to the presence of immunosuppressive innate immune cells, predominantly macrophages and monocytes^104,105^. Along this line, bladder cancer patients treated with the cyclooxygenase (COX) 1 and 2 inhibitor aspirin while undergoing intravesical immunotherapy benefited from better response rates^106^. In keeping with these results, it was previously shown that COX-2 driven inflammation stimulates the infiltration of immunosuppressive myeloid cells to the TME which in turn impairs responses to checkpoint inhibitors^107^. Thus, modulating the innate immune system, in particular macrophages will likely boost the anti-tumor effects of checkpoint inhibition^108^.

Our findings show that the zebrafish xenograft model has the potential to provide a real-time window with single-cell resolution to test and mechanistically understand new therapies targeting the innate immune system, in particular innate immunomodulatory drugs/vaccines. These new drugs/vaccines could be then combined with immune checkpoint therapies, to engage both arms of the immune system in the fight against cancer.

## MATERIALS AND METHODS

### Zebrafish husbandry

Zebrafish (*Danio rerio*) were handled and maintained according to the standard protocols of the European Animal Welfare Legislation, Directive 2010/63/EU (European Commission, 2016) and Champalimaud Fish Platform. All protocols were approved by the Champalimaud Animal Ethical Committee and Portuguese institutional organizations—ORBEA (Órgão de Bem-Estar e Ética Animal/Animal Welfare and Ethics Body) and DGAV (Direção Geral de Alimentação e Veterinária/Directorate General for Food and Veterinary).

Zebrafish, between 3 and 18 months old, were reared in 3.5L tanks at a density of 10 fish/L with females and males together. Rearing temperature was 28°C. Animals were kept in a light cycle of 14 hours (from 8 am until 10pm). Zebrafish were fed three times per day, artemia in the mornings, and powder (Sparos 400-600, Cat. No. U000001864 – Techniplast) in the afternoons and evenings.

### Zebrafish transgenic lines

According to the purpose of each experiment, different genetically modified zebrafish lines were used in this study: *Tg*(*mpx:eGFP*)^56^, *Tg(mpeg1:mCherry-F*)^109^, *Tg(csfr1a:GFP)*^71^ and *Tg(mpeg1:mCherry-F; tnfa:GFP-F*)^109^. All the transgenic and nacre fish were in the Tübingen background.

### Human cancer cell lines and culture

Human urothelial cancer RT112 (female) and J82 (male) cell lines were a kind gift from Dr. Mireia Castillo (Champalimaud Foundation, Portugal). Cell lines were authenticated by Small Tandem Repeat profiling using FTA cards (STABvida, Portugal) and were routinely mycoplasma tested. Both cell lines were kept and grown in Dulbecco’s modified Eagle medium (DMEM) high glucose (Biowest) and supplemented with 10% fetal bovine serum (FBS) (Sigma-Aldrich) and antibiotics (100 U mL^−1^ penicillin and 100 μg mL^−1^ streptomycin, Hyclone) in a humidified 5% CO_2_ atmosphere at 37 °C.

### Cell staining

Tumor cells were grown to 85-90% confluence in T-175 flasks, washed with Dulbecco’s phosphate-buffered saline (DPBS) 1X (Biowest) and detached enzymatically-TrypLE (Thermo Fisher). Cell suspension was collected to 15mL centrifuge tubes, spun down at 300 × *g*, for 4 min and resuspended in DPBS 1X. Cells were then stained in 1.5mL microcentrifuge tubes with lipophilic dyes—Vybrant CM-DiI (4 μl/mL in DPBS 1X) or Deep Red Cell Tracker (1 μl/mL in DPBS 1X, 10 mM stock) (Life Technologies), for 15 min at 37 °C, protected from light. Cells were washed by spinning down at 300 × *g*, for 5 min at 4 °C and resuspended in complete medium. Viability was assessed by the trypan blue exclusion method, and cell number was determined by hemocytometer counting. Cells were resuspended in complete medium to a final concentration of 0.5 × 10^6^ cells/μL.

### Bacterial strains

The recombinant Bacille Calmette-Guerin BCG*ΔureC::hly* (VPM1002)^47,48^, BCG:SSI pGFP^47,48^ and BCG:SSI pmCherry^47,48^ were provided by the Department of Immunology, Max-Planck Institute for Infection Biology-MPIIB, Germany. OncoTICE® (BCG Strain TICE® - Merck) was provided by the Urology Unit – Champalimaud Foundation, Portugal.

### Bacterial culture

Glycerol frozen bacteria were thawed on ice for ∼3-4 hours. Thawed bacteria were spun down at 3000 x *g* for 10 minutes and washed twice in PBS 1X. Pelleted bacteria were resuspended in 100μL of PBS 1X, seeded on Middlebrook 7H11 plates supplemented with 10% OACD and incubated at 37°C until colonies formed (∼4-5 weeks). Fluorescent BCG:SSI colonies were selected and grown in 5mL of liquid Middlebrook 7H9 broth supplemented with 10% ADC and Hygromycin 50μg/mL (Cat. No. H7772 – Sigma) in 50mL centrifuge tubes at 37°C, shaking at 50 RPM until high turbidity was reached. 1mL of highly concentrated bacterial culture was seeded into 9mL of Middlebrook 7H9 + 10% ADC + Hygromycin 50μg/mL in 30mL sterile bottles (Cat. No. 2019-0030 – Thermo Scientific) and incubated at 37°C, shaking at 50 RPM until they reached the OD600:0.8.

VPM1002 colonies were selected and grown in 5mL of liquid Middlebrook 7H9 broth supplemented with 10% ADC in 50mL centrifuge tubes at 37°C, shaking at 50 RPM until high turbidity was reached. 1mL of highly concentrated bacterial culture was seeded into 9mL of Middlebrook 7H9 + 10% ADC in 30mL sterile bottles and incubated at 37°C, shaking at 50 RPM until they reached the OD600:1.2.

Once the desired OD was reached, bacteria were spun down at 3000 x *g* for 10 minutes. Pelleted bacteria were then washed and resuspended in PBS 1X from which a sample was streaked in Middlebrook 7H11 plates for CFU enumeration. Bacteria were spun down again and resuspended in 10% glycerol/PBS solution, frozen in cryovials and stored at -80°C. In order to check for contamination, an aliquot of bacterial culture was streaked on LB agar plates at different time points of the protocol and incubated at 37°C.

OncoTICE® vials were resuspended in sterile sodium chloride 0.9% solution at the Day Hospital (Champalimaud Foundation) according to the manufacturer instructions (Merck: 1 vial/50mL saline solution). Remnants from resuspended vials were stored at 4°C and protected from light.

### Bacterial staining

OncoTICE® vials were spun down at 3000 x *g* for 10 minutes, the supernatant was carefully discarded and pelleted bacteria were resuspended in lipophilic dye solutions: Vybrant CM-DiI (4 μl/ml in PBS 1X) or Deep Red Cell Tracker (1 μl/ml in PBS 1X, 10 mM stock). Bacteria were then incubated at 37°C and 300 RPM for 30 minutes protected from light. Labelled bacteria were spun down at 3000 x *g* for 5 minutes, washed once with PBS 1X and resuspended to the desired concentration in PBS 1X.

### *In vitro* challenge with BCG

RT112 and J82 cells were seeded in 24-well plates previously lined with sterile coverslips and incubated in a humidified 5% CO_2_ atmosphere at 37 °C. Both cell lines were challenged on days 1 and 3 after seeding with either DPBS 1X (Control) or BCG 10X (OncoTICE® - 1-8 x 10^8^ CFUs). On day 4 after seeding, cell medium was removed, cells washed and fixed in 4% (v/v) FA for 10 minutes and immunofluorescent staining was immediately performed.

### Immunofluorescence staining for *in vitro* cultures

FA fixed cells were washed twice for 5 minutes with 500uL of PBS1x at room temperature (RT). Cells were permeabilized by incubation at RT with 0.1%Triton/PBS 1X for 25 minutes. Cells were blocked in 500uL of PBDX_GS (50mL PBS 1X, 0.5 gr bovine serum albumin-BSA, 0.5mL DMSO, 0.25mL 1%Triton and 0.75mL goat serum) for 1 hour at RT. Cells were then incubated in 30μL of primary antibody dilution (1:200 in PBDX_GS) on top of a sheet of parafilm, inside a humid chamber at 4°C overnight.

Next day, cells were washed three times with PBS1x for 5 minutes at RT. Cells were then incubated in 30μL of secondary antibody dilution (1:500 in PBDX_GS) on top of a sheet of parafilm, inside a humid chamber at 4°C overnight and protected from light. After incubation cells were washed twice in PBS 1X for 5 minutes at RT. Cells were incubated for 10 minutes in 50μg/mL DAPI (15:10000 dilution) protected from light. Cells were then washed twice in distilled water for 5 minutes at RT. Coverslips were then dried and mounted on microscope slides with the help of aqueous mounting medium. Slides were stored at 4°C protected from light.

### Zebrafish xenografts

On the injection day, hatched embryos were separated from unhatched eggs. Pronase 1X was added to the embryo medium to boost hatching. The embryos were anesthetized by incubation in Tricaine 1X for 5 minutes. ∼50 anesthetized embryos were transferred to an agar/agarose plate. The embryos were carefully aligned in the agar/agarose plate with the help of a hairpin-loop. Fluorescently labelled cancer cells were injected using a microinjection needle under a stereomicroscope (ZEISS Stemi 305) with a milli-pulse pressure injector (Applied Scientific Instrumentation – MPPI-3). The treated embryos were transferred to a clean standard petri dish with Tricaine 1X solution and left to rest for 10 minutes to give time for the wound to close. Treated embryos were then placed in fresh E3 medium and incubated at 34°C. At 1 day post-injection (dpi), zebrafish xenografts were screened for the presence or absence of tumoral masses in a fluorescent stereomicroscope (Zeiss Axio Zoom.V16). Xenografts with edema, cells in the yolk sac or cellular debris were discarded.

At 4dpi zebrafish xenografts were analyzed in a fluorescent stereomicroscope (Zeiss Axio Zoom.V16) and the clearance rate was quantified as follows:

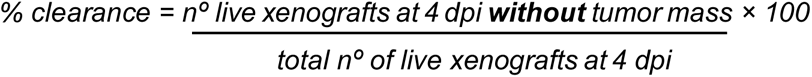

### Zebrafish macrophage ablation with clodronate liposomes

At 1dpi and 3dpi, xenografts were anesthetized by incubation in Tricaine 1X for 5 minutes. For the selective ablation of macrophages, ∼14nL of either liposome-encapsulated PBS (L-PBS) or liposome-encapsulated clodronate (L-clodronate) (Cat. No. CP-005-005 – Liposoma, 5mg/mL) were injected intratumorally at a 0.5X concentration using a microinjection needle under a stereomicroscope with a milli-pulse pressure injector. Treated xenografts were placed immediately in clean E3 medium and incubated at 34°C.

### Chemotherapy of zebrafish xenografts

At 1dpi, zebrafish were randomly distributed in control and treatment groups. Maximum tolerated concentration of drugs in zebrafish larvae was determined as previously described^22^. Zebrafish were then anesthetized by incubation in Tricaine 1X for 5 minutes and ∼14nL of either L-PBS, L-Clodronate, Mitomycin C(0.5mg/mL; Medac)/L-PBS or Mitomycin C/L-Clodronate were injected intratumorally and xenografts were placed immediately in clean E3 medium. This procedure was repeated at 3dpi. Throughout the experiment, xenografts were kept at 34°C and assessed daily. Xenografts were euthanized and fixed at 4dpi.

### BCG immunotherapy of zebrafish xenografts

At 1dpi, zebrafish were randomly distributed in control and treatment groups. BCG stock vials were thawed on ice, spun down at 3000 x *g* for 10 minutes and washed twice in PBS 1X. Bacteria were passed through a 25G needle to promote single cell dilution and resuspended in PBS 1X to a final concentration of 3-4 x 10^6^ CFU/mL. Xenografts were anesthetized with Tricaine 1X. ∼14nL of either L-PBS, L-Clodronate, BCG or BCG/L-Clodronate were injected intratumorally and xenografts were placed immediately in clean E3 medium. This procedure was repeated at 3dpi. Throughout the experiment, xenografts were kept at 34°C and assessed daily. Xenografts were euthanized and fixed at 4dpi.

### Single-cell light sheet live imaging and analysis of zebrafish xenografts

At 1 dpi, control or BCG/VPM1002 treated *Tg(*csf1ra:GFP*)^sh^*^377^ xenografts were left to rest in E3 medium for ∼5 minutes immediately after treatment. A single xenograft was then chosen and mounted in a capillary tube with 0.8% low melting agarose. The mounted xenograft was placed inside the chamber of a Zeiss Light Sheet Z.1 microscope, previously filled with Tricaine 0.75X in E3 medium without methylene blue at 34°C. Using a 20x objective lens and the Zen Blue software, the area of the tumor was delimited and z-stack images were acquired every 3 minutes within a 5μm interval. Xenografts were imaged for ∼15 hours and then euthanized.

Light sheet files were converted to HDF5/XML files using the BigDataViewer plugin from ImageJ/Fiji Software^110^. Randomly selected individual macrophages were manually tracked in 3D using the MaMut plugin from ImageJ/Fiji^111^. Motion analysis (max. distance traveled, total distance traveled and mean speed) was based on the TrackMate algorithms, ImageJ/Fiji^112^.

For the quantification of elongated macrophages, phagocytosis, macrophage touching and macrophage fusion, three maximum intensity projections (MIP) of each tumor were assessed. Namely, tumors were divided in thirds in relation to their Z plane and a MIP was created from each third. Then, each event was manually quantified along the 15 hours of imaging per MIP (∼900 images per tumor).

Data was exported as CSV files and statistical analysis was performed using GraphPad Prism 8.0 software.

### Immunofluorescence

Xenografts stored in 100% MetOH were rehydrated by a series of decreasing MetOH concentrations (75%, 50%, 25% MetOH/0.1% Triton PBS 1X). Xenografts were washed 4x for 5 minutes in 0.1% Triton/PBS 1X then washed 1x for 5 minutes in milliQ H_2_O. Xenografts were then incubated on ice cold acetone at -20°C for 7 minutes and washed 2x for 10 minutes in 0.1% Triton/PBS 1X. Then they were incubated at RT for 1 hour in PBDX_GS blocking buffer. PBDX_GS was removed and ∼40uL of primary antibody dilution was added (1:100 in PBDX_GS). Xenografts were incubated at RT for 1 hour and then overnight at 4°C.

Primary antibody was removed and xenografts were washed 2x for 10 minutes in 0.1% Triton/PBS 1X. Then, they were washed 4x for 30min in 0.05% Tween/PBS 1X. 0.05% Tween/PBS 1X was removed and ∼40uL of secondary antibody dilution (1:200 in PBDX_GS) + DAPI (50μg/mL) was added. Xenografts were incubated at RT for 1 hour and then overnight at 4°C.

Secondary antibody dilution was removed and xenografts were washed 4x for 15min in 0.05% Tween/PBS 1X. Xenografts were fixed in 4% FA for 20 minutes and washed 1x in 0.05% Tween/PBS 1X for 10 minutes. Xenografts were then mounted in Mowiol aqueous mounting medium (Cat. No. 81381 – Sigma) between two coverslips to allow for double side microscope acquisition.

### Confocal imaging and analysis of zebrafish xenografts

Mounted xenografts were imaged using an inverted LSM 710 confocal microscope (Zeiss) with Zen software. Tumors were imaged with a 25x immersion objective lens using the z-stack function with an interval of 5μm between slides. The number of cells was manually assessed with the cell counter plugin from ImageJ/Fiji. To assess tumor size, three representative slides of the tumor, from the top (Zfirst), middle (Zmiddle), and bottom (Zlast), per z-stack per xenograft were analyzed and a proxy of the total cell number (DAPI nuclei) was estimated as follows:

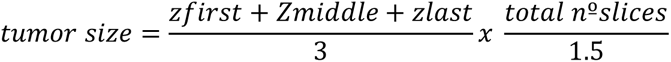

The 1.5 correction number was estimated to human cells that have a nucleus with an average of 10–12μm of diameter. The number of activated caspase-3 positive cells, macrophages, neutrophils, TNFα positive/negative macrophages was individually quantified in every slide along the tumor. To get the percentage of each population, the obtained number was divided by its corresponding tumor size.

### Histopathology

Fish were euthanized, fixed in 4% PFA and longitudinally embedded in paraffin. 4μm serial sections were cut and stained with Hematoxylin and Eosin, and Ziehl-Neelsen (Cat. No. R0276 – Liofilchem). Tissue sections were examined by a pathologist from the Champalimaud Foundation Histopathology platform in an Axioscope 5 microscope (Zeiss) and microphotographs captured in an Axiocam 208 color camera (Zeiss).

### Whole-mount *in situ* hybridization

4dpi zebrafish xenografts were collected and fixed in 4% formaldehyde at 4°C overnight, dehydrated through a methanol series and stored in 100% methanol at –20°C. Whole-mount *in situ* hybridizations were performed as described^113^ with minor modifications (hybridization temperature 65°C), using digoxigenin (DIG) labelled antisense RNA probes for l-plastin and c-myb. Staining reaction was performed using BMP-Purple (Roche). Zebrafish larvae xenografts were photographed using a Zeiss SteREO Discovery.V8 coupled to a Zeiss AxioCam Icc 3 Camera.

### Statistical analysis

Statistical analysis was performed using the GraphPad Prism 8.0 software. All data sets were challenged by D’Agostino & Pearson and Shapiro–Wilk normality tests. In general, data sets with a Gaussian distribution were analyzed by parametric unpaired *t* test and data sets that did not pass the normality tests were analyzed by nonparametric unpaired Mann–Whitney test. Clearance data sets were analyzed using Fisher’s exact test. All were two-sided tests with a confidence interval of 95%. Differences were considered significant at *P* < 0.05 and statistical output was represented as follows: non-significant (ns) ≥0.05, *<0.05, **<0.01, ***<0.001, ****<0.0001. Bars indicate the results as AVG ± standard deviation of the mean (STDEV).

## ACKNOWLEDGEMENTS

We thank the Champalimaud Foundation, Congento (LISBOA-01-0145-FEDER-022170, co-financed by FCT/Lisboa2020) and FCT-PTDC/MEC-ONC/31627/2017 for funding. FCT fellowships for Mayra Martínez-López (PD/BD/138203/2018), and Catia Rebelo de Almeida (2021/08619/BD). We are grateful to all members of Fior Lab for their support and critical discussion; Dr. Mireia Castillo for the procurement of cell lines; the CF Fish Platform (Catarina Certal, Joana Monteiro et al.) for excellent animal care; the CF Glass Wash and Media Preparation Platform (Maria Vito); the CF Histology Platform (Tania Carvalho et al.); the CF Advanced BioImaging and BioOptics Experimental Platform (Davide Accardi and Anna Pezzarossa); and the CF Molecular and Transgenic Tools (Catarina Certal, Ana Cunha and Raquel Tomás) for their technical support. We are also grateful to Dr. Gopinath Krishnamoorthy, Dr. Martin Rao and Dr. Pedro Moura Alves for their technical support and advice regarding BCG culture and experiments; Dr. Iván Moya for critical discussion of the manuscript, and to the zebrafish community for their generosity in sharing fish strains (Stephen Renshaw, Farida Djouad, and Zilong Wen).

## DATA AVAILABILITY

The data supporting the findings of this study are available within the article and its supplementary materials. Any additional information required to reanalyze the data reported in this paper is available from the corresponding author upon request.

## AUTHOR CONTRIBUTIONS

M.M-L. planned and performed all the experiments, analyzed the data, wrote and revised the manuscript. C.R.dA. performed the experiments, supervised M.F., analyzed the data and revised the manuscript. M.F. performed the TNF inhibition experiments, analyzed the data and revised the manuscript. R.V.M. performed the in situ experiments, analyzed the data and revised the manuscript. S.H.E.K. provided the live BCG strains provided feedback and revised the manuscript. R.F. supervised all research, planned and performed experiments, wrote and revised the manuscript.

---

**Video 1. Macrophage kinetics of control NMIBC-RT112 zebrafish xenografts 1dpi.** Maximum intensity projection of the tumor. Each colored line represents the path a single macrophage followed in a 15-hour time lapse. Images of the tumor were acquired in stacks of 5 µm in the Z plain every 3 minutes. Tracking was made using the MaMut plugin from ImageJ/Fiji.

**Video 2. Macrophage kinetics of BCG treated NMIBC-RT112 zebrafish xenografts 1dpi.** Maximum intensity projection of the tumor. Each colored line represents the path a single macrophage followed in a 15-hour time lapse. Images of the tumor were acquired in stacks of 5 µm in the Z plain every 3 minutes. Tracking was made using the MaMut plugin from ImageJ/Fiji.

**Video 3. Macrophage kinetics of VPM1002 treated NMIBC-RT112 zebrafish xenografts 1dpi.** Maximum intensity projection of the tumor. Each colored line represents the path a single macrophage followed in a 15-hour time lapse. Images of the tumor were acquired in stacks of 5 µm in the Z plain every 3 minutes. Tracking was made using the MaMut plugin from ImageJ/Fiji.

**Video 4. Macrophage touching in the TME of NMIBC-RT112 zebrafish xenografts.** Representative video showing macrophages (labelled in green) phagocyting cancer cells (labelled in magenta) and actively touching their cell membranes within the tumor microenvironment of a 1dpi NMIBC-RT112 zebrafish xenograft.

**Video 5. Macrophage fusion-like events in the TME of NMIBC-RT112 zebrafish xenografts.** Representative video showing macrophages (labelled in green) phagocyting cancer cells (labelled in magenta) and joining their cell membranes within the tumor microenvironment of a 1dpi NMIBC-RT112 zebrafish xenograft.

**Video 6. Dendritic-like cells in the TME of NMIBC-RT112 zebrafish xenografts.** Representative video showing macrophages (labelled in green) and cancer cells (labelled in magenta) within the tumor microenvironment of a 1dpi bladder cancer xenograft. Dendritic-like cells with no phagocytic behavior can be seen actively interacting with their surrounding macrophages.

## SUPPLEMENTARY FIGURE LEGENDS

**Supplementary figure 1.**
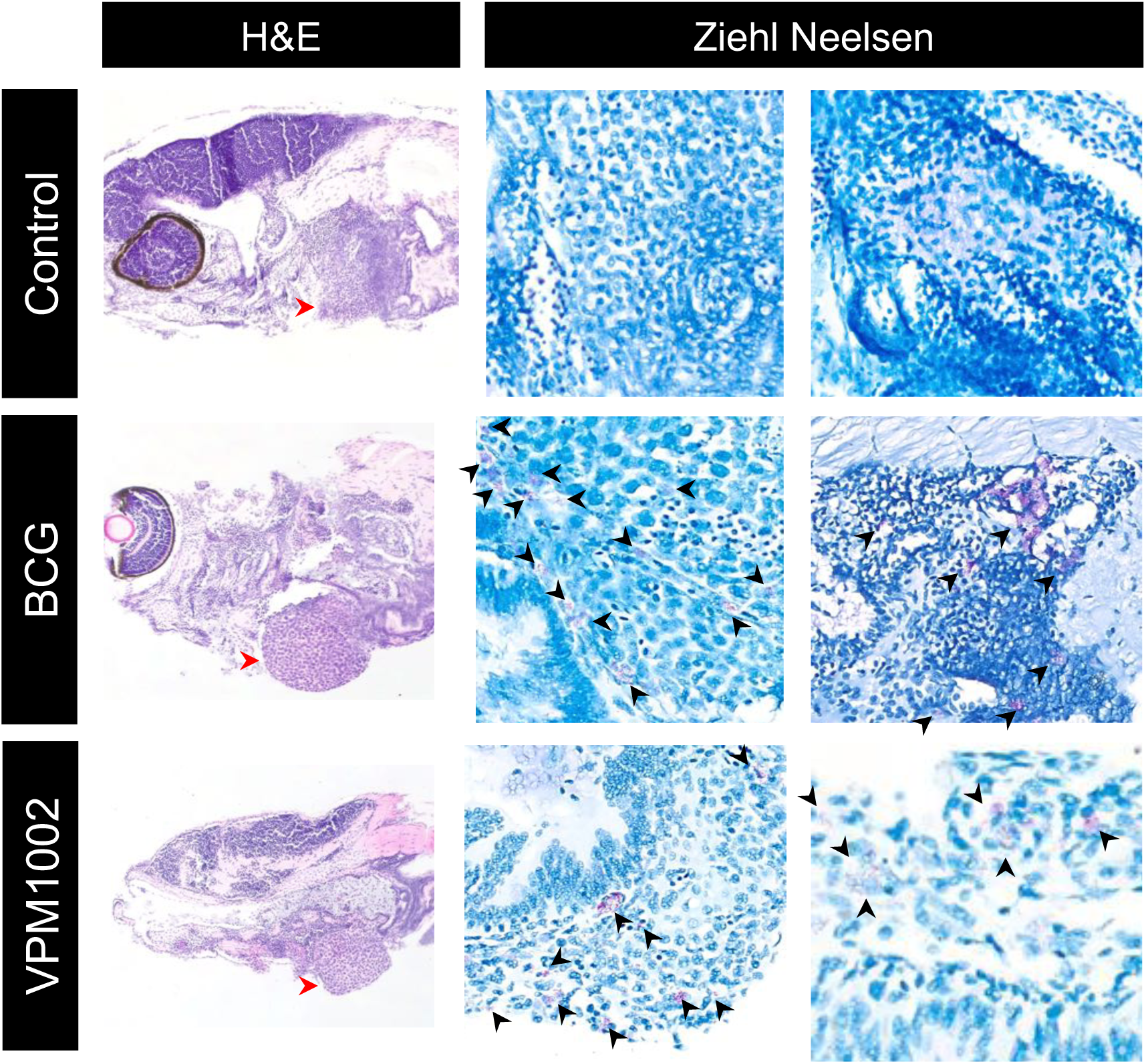
A zebrafish xenograft model for BCG immunotherapy in bladder cancer. Representative microphotographs of zebrafish xenografts, stained with Hematoxylin and Eosin (first column, red arrow heads point to the tumor) and with Ziehl Neelsen (second and third column). Acid-fast bacilli, staining bright red with Ziehl Neelsen (black arrow heads), are seen within some of the tumors, inside macrophages, extracellularly and, more rarely, inside tumor cells.

**Supplementary figure 2.**
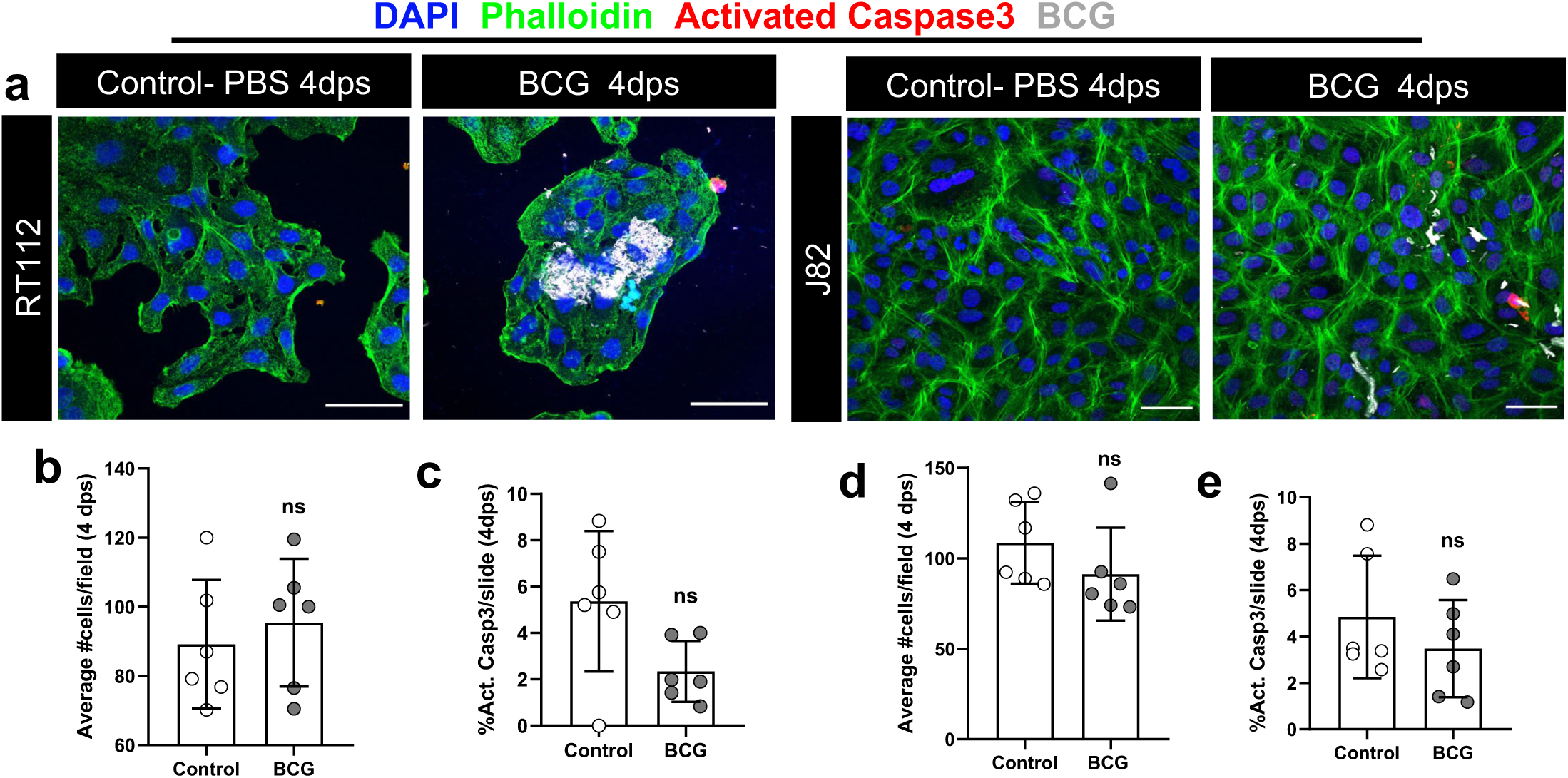
NMIBC-RT112 and MIBC-J82 cell lines are not susceptible to BCG *in vitro*. **a)** Representative confocal images of NMIBC-RT112 and MIBC-J82 cells stained for the actin filaments marker phalloidin (green), apoptosis marker activated caspase 3 (red), BCG (white) and DAPI nuclei counterstaining. **b)** Quantification of the mean absolute number of cells per field in control and treated NMIBC-RT112 cells at 4dps. **c)** Quantification of the percentage of activated caspase 3 cells per field in control and treated NMIBC-RT112 cells at 4dps. **d)** Quantification of the mean absolute number of cells per field in control and treated MIBC-J82 cells at 4dps. **e)** Quantification of the percentage of activated caspase 3 cells per field in control and treated MIBC-J82 cells at 4dps. Bars indicate the results as AVG ± STDEV and each dot represents one quantified well. Data pooled from 2 independent experiments. Scale bar: 50 µm. dps: days post-seeding.

**Supplementary figure 3.**
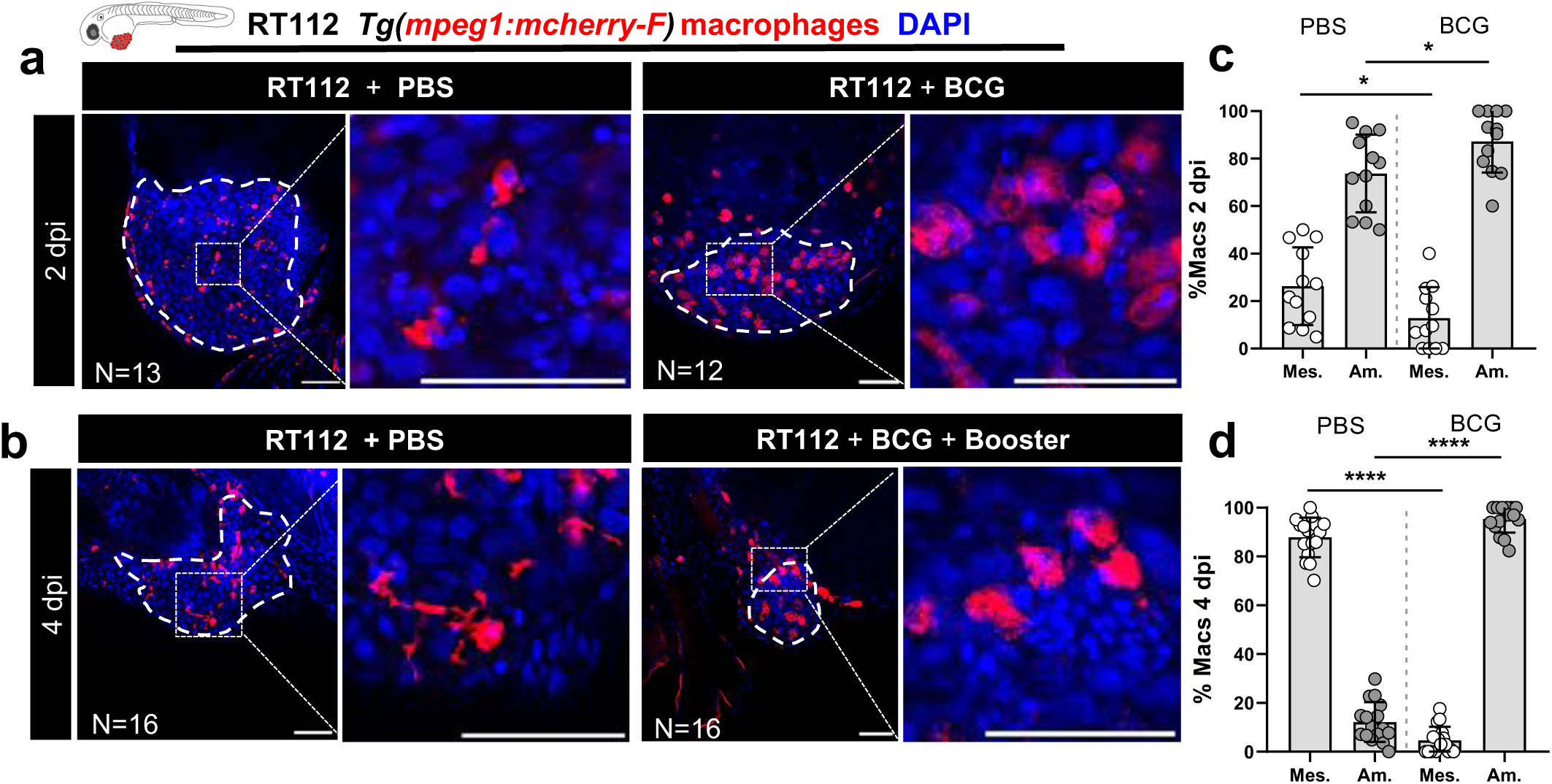
BCG treated xenografts comprise more macrophages with ameboidal morphology. **a and b)** Representative confocal images of infiltrating macrophages (red) in NMIBC-RT112 control and BCG-treated xenografts at 2 and 4dpi. **c and d)** Quantification of the percentage of infiltrating macrophages with either a mesenchymal or ameboidal morphology in NMIBC-RT112 control and BCG-treated xenografts at 2dpi (mesenchymal *P=0.0370, ameboidal *P=0.0370) and 4dpi (****P<0.0001). Bars indicate the results as AVG ± STDEV and each dot represents one xenograft pooled from 2 independent experiments. White dashes outline the tumor. All images are anterior to the left, posterior to right, dorsal up and ventral down. Scale bar: 50 µm. dpi: days post-injection.

**Supplementary figure 4.**
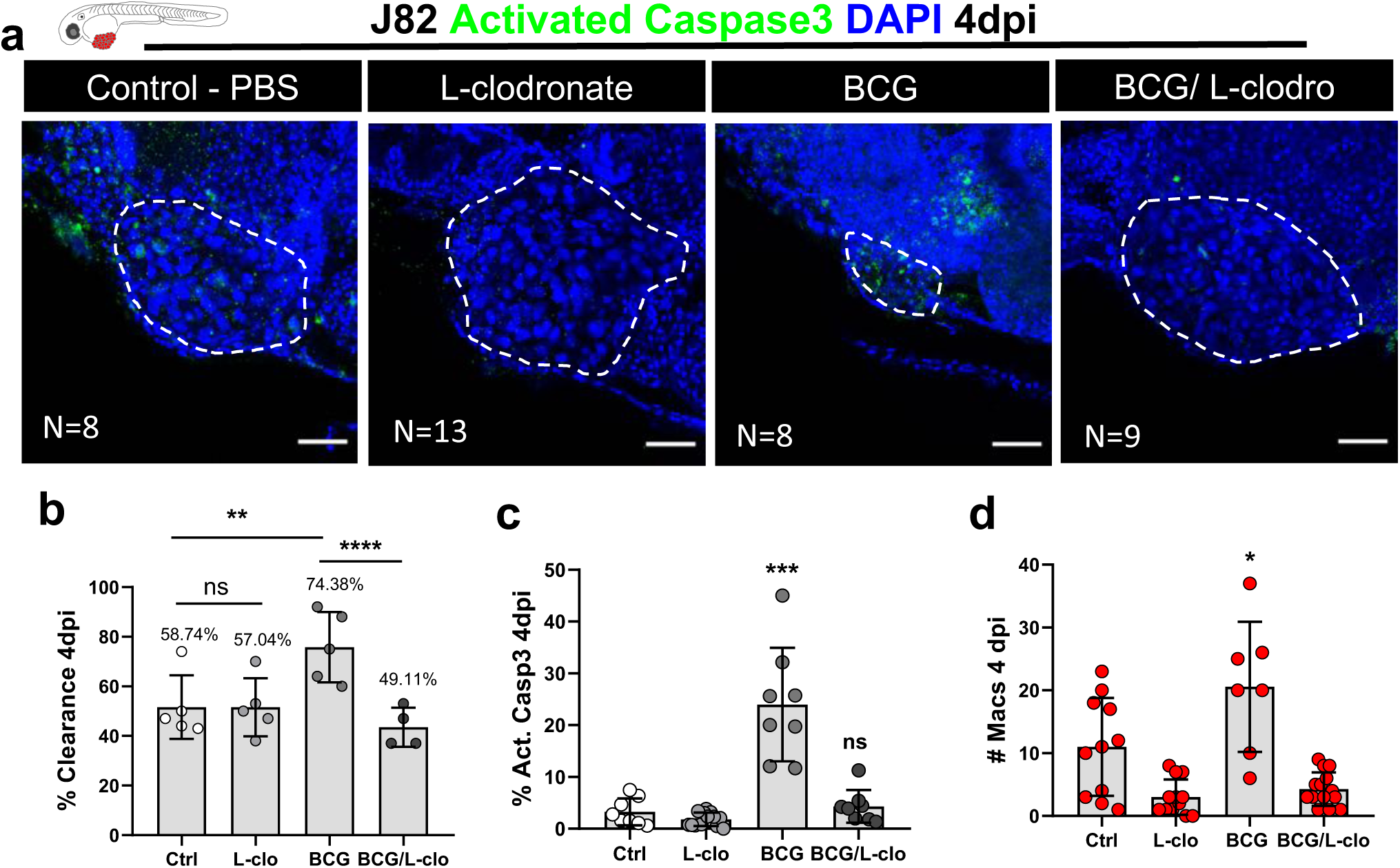
Macrophages are essential for susceptibility to BCG immunotherapy of J82 zebrafish bladder cancer xenografts. **a)** Representative confocal images of MIBC-J82 xenografts stained for the apoptosis marker activated caspase 3 (green) with DAPI nuclei counterstaining in BCG/L-clodronate experiments at 4dpi. **b)** Quantification of the percentage of clearance in BCG/L-clodronate experiments at 4dpi (**P=0.0091, ****P<0.0001). Bars indicate the results as AVG ± STDEV and each dot represents a full round of injections in which N= # of xenografts without tumor at 4dpi/ total number of xenografts at 4dpi**. c)** Quantification of the percentage of apoptosis/activated caspase3 positive cells in BCG/L-clodronate experiments at 4dpi (***P=0.0002). **d)** Quantification of the absolute numbers of infiltrating macrophages in BCG/L-clodronate experiments (*P=0.0461). Bars indicate the results as AVG ± STDEV and each dot represents one xenograft pooled from 3 independent experiments. All images are anterior to the left, posterior to right, dorsal up and ventral down. White dashes outline the tumor. Scale bar: 50 µm. dpi: days post-injection.

**Supplementary figure 5.**
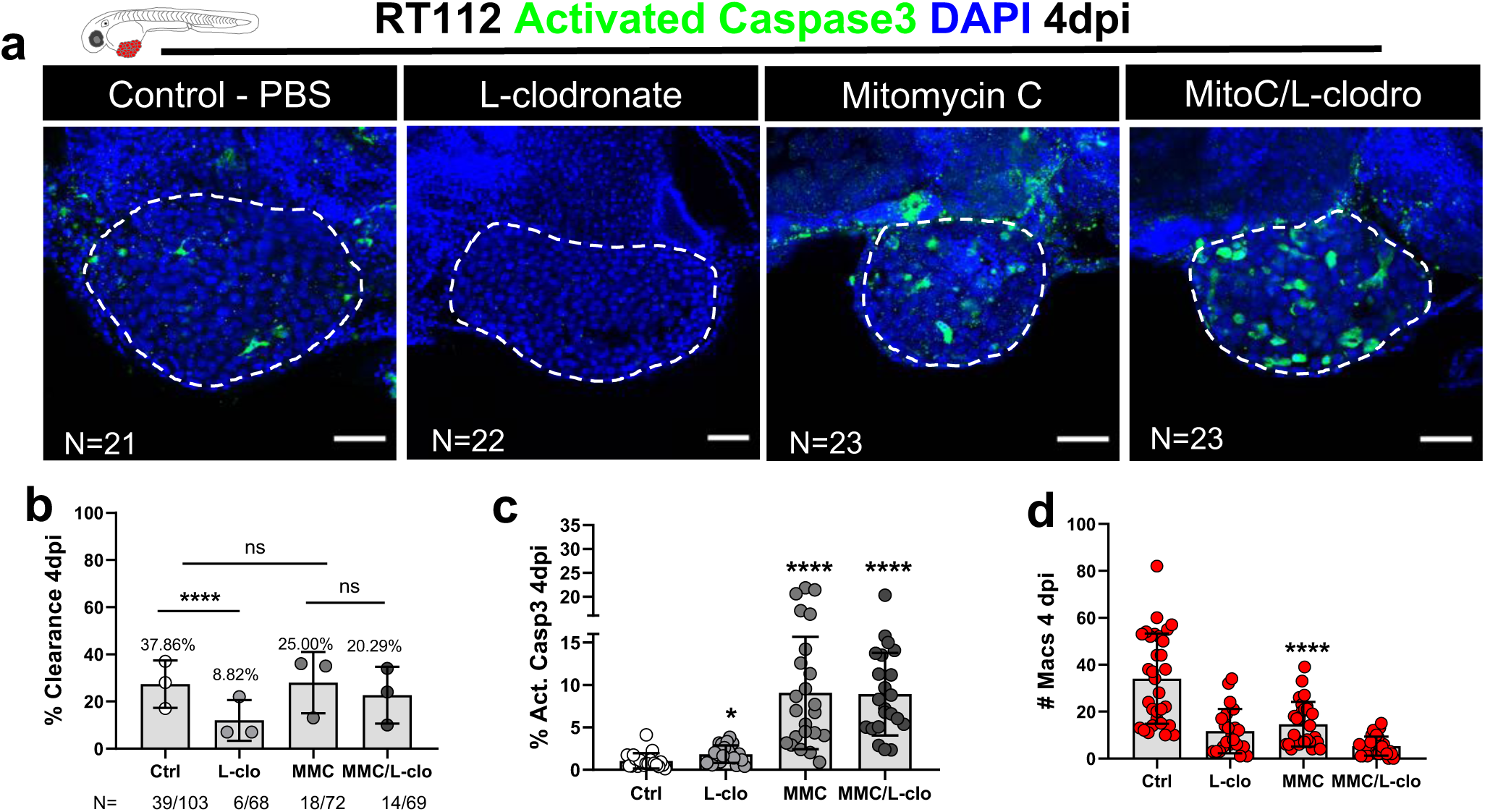
Cytotoxic effects of Mitomycin C in zebrafish bladder cancer xenografts are not mediated by macrophages. **a)** Representative confocal images of NMIBC-RT112 xenografts stained for the apoptosis marker activated caspase 3 (green) and DAPI nuclei counterstaining in MMC/L-clodronate experiments at 4dpi. **b)** Quantification of the percentage of clearance in MMC/L-clodronate experiments at 4dpi (****P<0.0001). Bars indicate the results as AVG ± standard deviation of the mean (STDEV) and each dot represents a full round of injections in which N= # of xenografts without tumor at 4dpi/ total number of xenografts at 4dpi**. c)** Quantification of the percentage of apoptosis/activated caspase3 positive cells in MMC/L-clodronate experiments at 4dpi (*P=0.0127, ****P<0.0001). **d)** Quantification of the absolute numbers of infiltrating macrophages in MMC/L-clodronate experiments (****P<0.0001). Bars indicate the results as AVG ± STDEV and each dot represents one xenograft pooled from 3 independent experiments. White dashes outline the tumor. All images are anterior to the left, posterior to right, dorsal up and ventral down. Scale bar: 50 µm. dpi: days post-injection. MMC: Mitomycin C.

**Supplementary figure 6.**
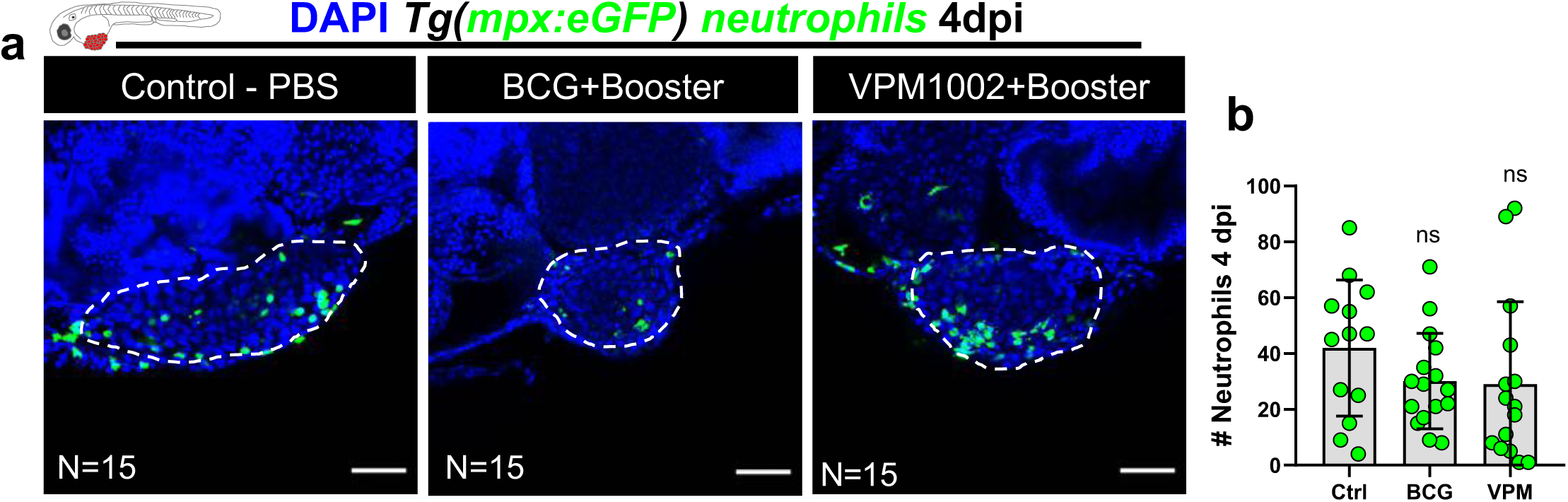
BCG treatment has no significant effects in neutrophil infiltration at 4dpi. **a)** Representative confocal images of neutrophils (green) in NMIBC-RT112 control and BCG- or VPM1002-treated xenografts at 4dpi. **b)** Quantification of the absolute numbers of infiltrating neutrophils at 4dpi. Bars indicate the results as AVG ± STDEV and each dot represents one xenograft pooled from 2 independent experiments. White dashes outline the tumor. All images are anterior to the left, posterior to right, dorsal up and ventral down. Scale bar: 50 µm. dpi: days post-injection.

**Supplementary figure 7.**
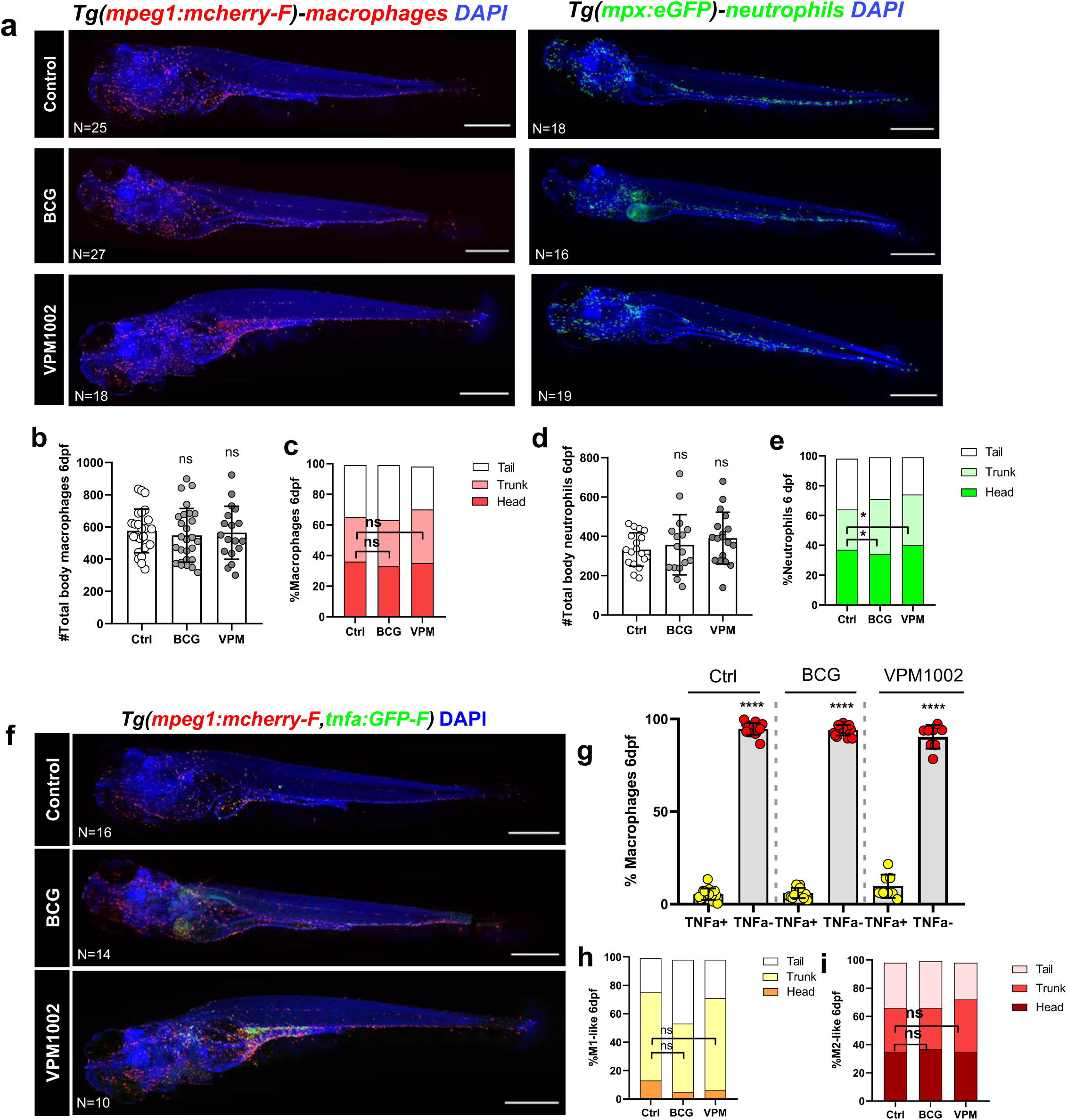
BCG treatment has no significant effects in neutrophil nor macrophage distribution and polarization in zebrafish larvae. **a,f)** Representative full body confocal images of macrophages (red), neutrophils (green), TNFα expression (green) and macrophages (red) in 4dpi/6dpf control and BCG or VPM1002 treated larvae. **b,d)** Quantification of the absolute number of total body macrophages and neutrophils. **c,e)** Distribution of macrophages and neutrophils in the larvae’s body (Ctrl vs BCG *P=0.0116, Ctrl vs VPM1002 *P=0.0116) . **g)** Quantification of the percentage of TNFα positive/negative macrophages in the larvae’s body (****P<0.0001). **h,i)** Distribution of TNFα positive/negative macrophages in the larvae’s body. Bars indicate the results as AVG ± STDEV and each dot represents one larva pooled from 2 independent experiments. All images are anterior to the left, posterior to right, dorsal up and ventral down. Scale bar: 200 µm. dpi: days post-injection. dpf: days post-fertilization.

**Supplementary figure 8.**
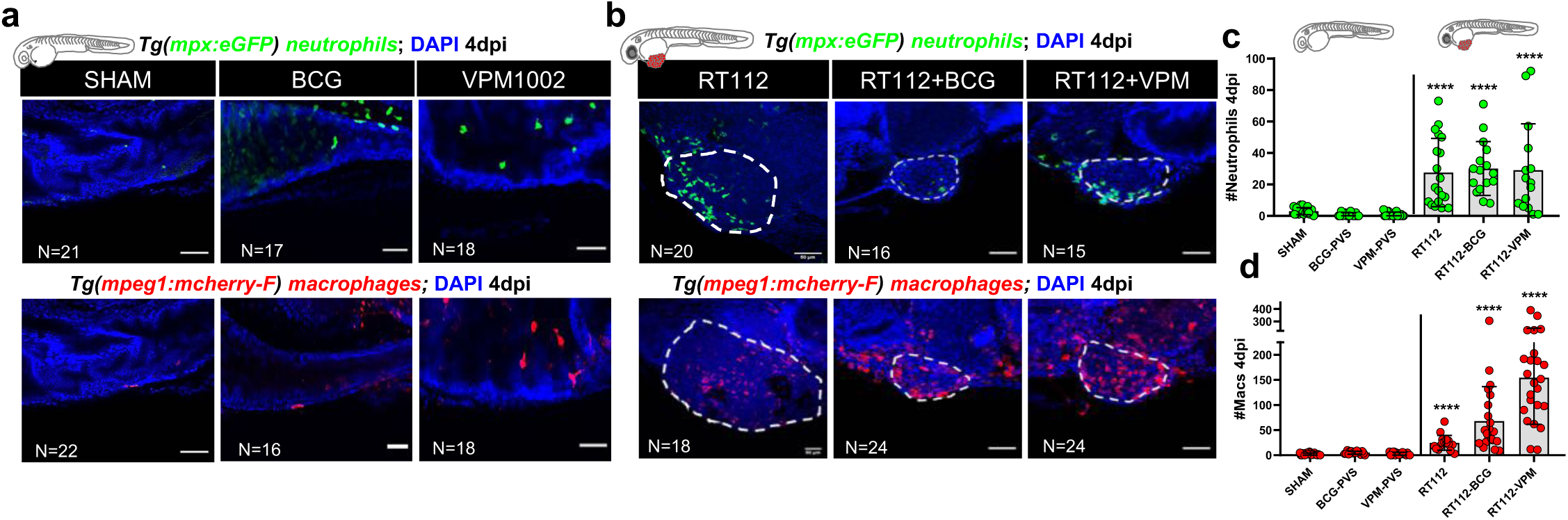
Bladder cancer cells are required for the recruitment of neutrophils and macrophages to the PVS in response to BCG immunotherapy. **a and b)** Representative confocal images of neutrophils (green) and macrophages (red) in the perivitelline space (PVS) of zebrafish larvae at 4dpi/6dpf. **c)** Quantification of the absolute numbers of neutrophils in the PVS of zebrafish larvae at 4dpi/6dpf (****, P<0.0001). **d)** Quantification of the absolute numbers of macrophages in the PVS of zebrafish larvae at 4dpi/6dpf (****, P<0.0001). Neutrophil and macrophage data sets were compared against their corresponding SHAM control. Bars indicate the results as AVG ± STDEV and each dot represents one xenograft pooled from 3 independent experiments. White dashes outline the tumor. All images are anterior to the left, posterior to right, dorsal up and ventral down. Scale bar: 500 µm. Scale bar: 50 µm. dpi: days post-injection; dpf: days post-fertilization.

**Supplementary figure 9.**
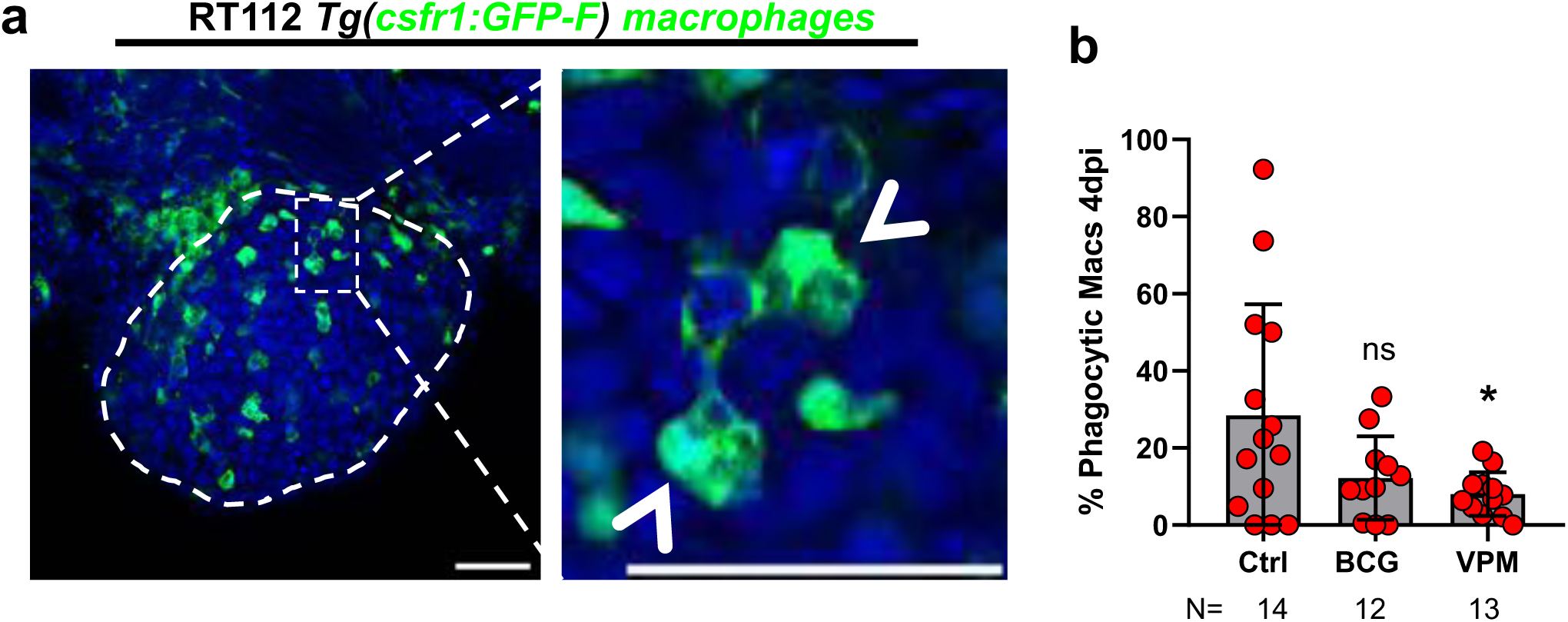
Not all infiltrating macrophages engage in phagocytosis within the TME. **a)** Representative confocal images of macrophages (green) within the TME. **b)** Quantification of the percentage of macrophages engaged in phagocytosis within the TME (*P=0.0208). Bars indicate the results as AVG ± STDEV and each dot represents one xenograft pooled from 2 independent experiments. White dashes outline the tumor. White arrow heads show phagocytic macrophages. All images are anterior to the left, posterior to right, dorsal up and ventral down. Scale bar: 50 µm. dpi: days post-injection.

